# The major role of the REL2/NF-*κ*B pathway in the regulation of midgut bacterial homeostasis in the malaria vector *Anopheles gambiae*

**DOI:** 10.1101/2025.03.14.643338

**Authors:** Suzana Zakovíc, Galo E. Rivera, Remmora Gomaid, Cristina Graham Martinez, Christine Kappler, Eric Marois, Elena A. Levashina

## Abstract

Multicellular organisms harbor diverse microbial communities that play essential roles in host physiology. While often beneficial, these interactions require tight regulation to prevent dysbiosis and disease. This study examines the tissue-specific immune responses of mosquitoes to blood feeding and *Plasmodium falciparum* infection in *Anopheles* females. We demonstrate that REL2 signaling regulates antimicrobial peptide (AMP) expression and shapes midgut bacterial composition post-blood meal. Loss of REL2 leads to midgut dysbiosis, characterized by the overgrowth of *Serratia* spp., and mosquito lethality within a day of feeding. Interestingly, *Serratia*-induced dysbiosis also reduces *P. falciparum* prevalence in surviving mosquitoes. Our findings highlight the critical role of immune system in maintaining midgut bacterial homeostasis and uncover complex interactions between mosquito immunity, gut microbiota, and malaria parasites.

**Graphical abstract:** 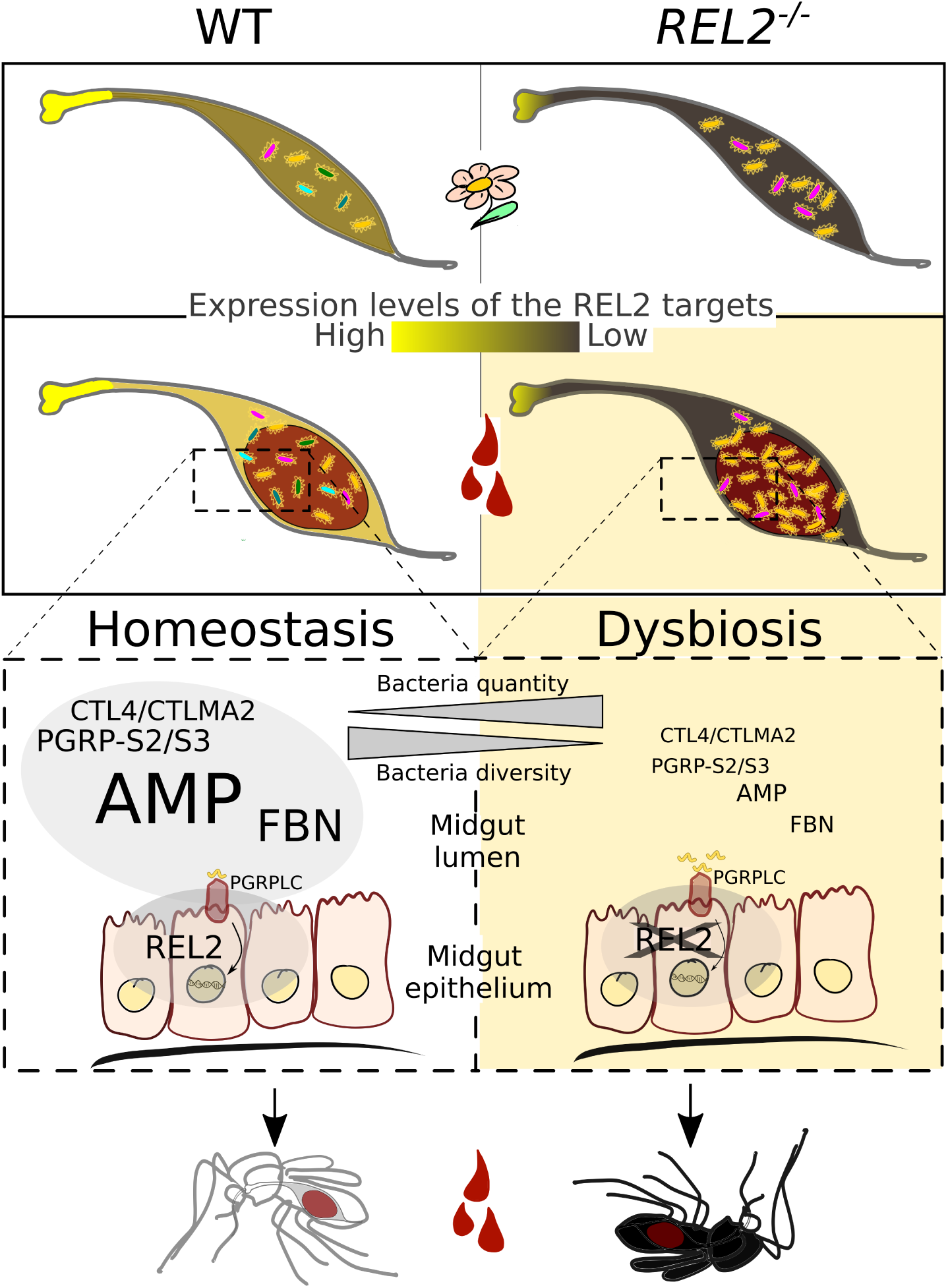

## Introduction

All multicellular organisms are inhabited by microbes that critically contribute to their physiology. While most microbes are harmless or beneficial to their hosts, such homeostatic interactions must be tightly controlled as even small perturbations in the system could lead to dysbiosis and disease.

The immune system plays a crucial role in maintaining homeostatic interactions with microbial communities at epithelial surfaces in mammals, a topic currently under intensive investigation. In insects, most research has focused on the microbiomes of the digestive tract. While beetles and bees harbor stable, well-defined microbial communities within distinct gut compartments^1,2^, mosquito midguts feature a more diverse and dynamic microbiome ^3,4^. Importantly, disruptions to these characteristic microbial communities by opportunistic environmental microorganisms can lead to pathogenic situations. However, how the immune system responds to nutritional stressors and other perturbations in gut microbial communities remains poorly understood. Studies in *Drosophila melanogaster* have been instrumental in elucidating insect antibacterial responses, revealing that a healthy immune response requires a balance between microbial sensing, signal transduction, and effector function. Recognition of conserved microbial structures by a diverse array of molecules triggers both humoral and cellular immune responses, shaping interactions between the host and its microbiota ^5^.

*Drosophila*’s humoral immunity is characterized by the induction of small cationic peptides with potent antimicrobial activity, collectively called antimicrobial peptides (AMPs). Microbial sensing by peptidoglycan recognition receptors (PGRPs) activates immune signaling pathways that culminates in AMP expression. Cellular immune responses, such as phagocytosis and encapsulation, are activated through scavenger receptors, thioester containing proteins (TEPs) and lectins. The recognition of microbial molecules can also trigger a series of extracellular proteolytic cascades leading to melanization, or deposition of melanin at the site of infection or injury, which is mediated by both cellular components (hemocytes) and humoral factors, including CLIP-serine proteases and prophenoloxidases.

The Toll and IMD NF-*κ*B-like pathways are key signaling modules in *Drosophila* that mediate immune responses triggered by bacteria and fungi^6^. The Toll pathway is activated mainly by l-lysine-type (Lys-type) peptidoglycans (PGNs) of Gram-positive bacteria and by *β*-glucans of fungi through binding to circulating receptors (PeptidoGlycan-Recognition Protein – Short A (PGRP-SA), Gram-Negative bacteria Binding Protein 1 (GNBP1) and GNBP3), leading to activation of the transcription factor Dif and expression of such AMPs as Drosomycins, Metchnikowin, and Bomanins ^7–9^. The IMD pathway is induced by sensing of meso-diaminopimelic acid-type (Dap-type) PGNs of Gram-negative bacteria by the membrane-bound PGRP-LC and intracellular PGRP-LE receptors, culminating in the activation of transcription factor Relish, and expression of Attacin, Cecropin, Defensin, Diptericin and Drosocin. IMD activity in the fruit fly’s midgut is essential for post-exposure control of pathogenic bacteria and maintenance of a healthy microbiome ^10–14^. In systemic infections, when bacteria are introduced into body cavity by pricking or injections and spread through hemolymph, the main responding tissue is the fat body, analogue of the mammalian liver, and the blood cells, hemocytes^15,16^. However, the physiological functions of the IMD pathway and AMPs in the midgut remain unknown.

The life style and microbial exposure of *Anopheles gambiae* mosquitoes, a major vector of the human malaria parasite *Plasmodium falciparum* in Africa, differ significantly from those of *Drosophila*. Adult *Anopheles* mosquitoes acquire microbes through feeding on water and plant nectars. In addition to nectar, females require a mammalian blood meal, which is essential for ovarian development and egg production. This nutrient-rich diet also fuels rapid proliferation of midgut bacteria, peaking one day after feeding^17–20^. These immune and physiological challenges trigger extensive transcriptional changes and tissue remodeling, including the synthesis by the anterior midgut cells of a protective chitinous structure enriched with glycoproteins that encapsulates the blood meal called the peritrophic matrix ^21,22^. Blood feeding also exposes female mosquitoes to *Plasmodium* infection. Once in the midgut lumen, malaria parasites undergo sexual reproduction, producing ookinetes - the motile stage that invades the midgut epithelium. This invasion coincides with the peak of bacteria proliferation in the posterior midgut. Successful ookinetes traverse the epithelium and convert into oocysts on the basal side of the midgut, where parasite development continues until the formation of sporozoites.

The regulation of immune responses in *Anopheles* mosquitoes differs from that in *Drosophila* ^23^. For example, mosquitoes lack the intracellular receptor PGRP-LE, while PGRP-LC has been shown to bind PGNs from both Gram-positive and Gram-negative bacteria^24^. Additionally, mosquitoes do not produce Diptericins, Drosocins or any AMPs regulated by the Toll pathway. Instead, their major AMPs include the conserved Defensins (DEF1-5), Cecropins (CEC1-4), Attacin and the mosquito-specific Gambicin (GAMB)^25^.

The highest expression levels of AMP transcripts (*CEC1-4*, *DEF1*, and *GAMB*) are found in the most anterior part of the *Anopheles* midgut, known as the cardia or proventriculus ^26,27^. This region likely acts as the first protective barrier, limiting pathogen entry into the midgut. Functionally, the cardia serves as a sphincter, regulating the passage of food from the foregut and crop to the narrow, esophagus-like anterior midgut, which then leads to the larger posterior midgut – functionally analogous to the mammalian stomach. Enzymatic and transcriptomic data suggest that the anterior midgut is involved in processing sugars and proteins, while the posterior midgut specializes in digesting blood and absorbing lipids, with enzymatic activities increasing after blood meal intake^26–28^. Currently, a tissue-specific gene expression atlas for the midgut tissues of *Anopheles* mosquitoes, including the cardia, anterior, and posterior midgut, is only available for unfed mosquitoes. In contrast, transcriptional changes in midgut regions following a blood meal have been exclusively documented in *Aedes aegypti* ^27^. A notable difference between unfed *Ae. aegypti* and *A. gambiae* in the posterior midgut is the high expression levels of AMPs and the low expression levels of digestive serine endopeptidases in *A. gambiae* ^27^. This suggests potentially distinct immune and digestive strategies between the two species. However, it remains unknown whether these differences persist after blood feeding, highlighting a significant knowledge gap in our understanding of the post-blood meal transcriptional landscape in *Anopheles*.

The regulation of immune responses and AMP function in *Anopheles* remains only partially understood. RNA interference (RNAi) studies aimed at identifying effector genes of the Toll/REL1 and IMD/REL2 pathways suggest that both contribute to the transcriptional regulation of numerous immune genes ^29,30^. However, an earlier *in vitro* study reported that REL2 specifically regulates expression of *CEC1, CEC3, GAMB* and *Leucine-Rich-Repeat Immune1 (LRIM1)* genes^31^. Functional studies of the REL2 pathway yielded contradictory results regarding its role in antibacterial and antiparasitic responses. For example, RNAi-mediated silencing of a negative regulator or transgenic overexpression of the REL2 transcription factor has produced varied outcomes, ranging from no detectable effect to complete mosquito resistance to *Plasmodium* infections ^30,32,33^. Conversely, down-regulation of REL2 pathway only mildly increased mosquito susceptibility to parasites^24,34,35^. However, it significantly promoted bacterial overgrowth in the midgut^20,24^, and increased mosquito mortality in systemic bacterial infections ^24,31^. Evidence also suggests that bacteria may contribute to mosquito immune responses against parasites or directly inhibit *Plasmodium* development in the midgut. Indeed, antibiotic treatments have been shown to increase parasite loads ^20,24,36,37^. Despite these findings, the interplay between the REL2 pathway, midgut bacteria, and *Plasmodium* infections remains poorly understood.

Our incomplete understanding of these interactions can be attributed to the challenge of integrating results from diverse approaches used in the studies. Previous research on gene targets of immune pathways has often been conducted at the whole-organism level using RNAi-mediated knockdowns, which can exhibit variable tissue-specific efficiency ^29,30^. In contrast, tissue-specific studies have identified significant effects of *REL2* overexpression on metabolic and regulatory genes, while detecting minimal impact on immune genes. These findings suggest the potential activation of compensatory mechanisms that mitigate signaling overload caused by REL2 misexpression^33,38^. Furthermore, antibacterial responses have typically been examined under non-physiological conditions, such as thoracic injections or oral feeding with excessive doses of non-commensal bacteria species.

The *Anopheles* mosquito’s lifestyle – characterized by repeated blood feedings that drive bursts of bacterial proliferation and *Plasmodium* infection - provides an ideal model for studying immune responses in a physiological context. In this study, we investigated the tissue-specific responses of *Anopheles* to blood feeding and *P. falciparum* infection in both wild-type and REL2-deficient mosquitoes, focusing on key immune tissues: blood cells and fat body, the cardia with anterior midgut, and the posterior midgut. Our findings show that REL2 plays a critical role in regulating AMP expression in both the anterior and posterior midgut 24 hours after blood feeding. Moreover, microbiome sequencing indicated that REL2 function at this stage shapes bacterial composition and is essential for female survival. Notably, we demonstrate that the inhibition of mosquito colonization by *P. falciparum* is caused by proliferation of *Serratia* spp. in the midgut of *REL2* mutants. These results identify REL2 as a key factor in maintaining midgut bacterial homeostasis and offer new insights into the complex interactions between mosquito immunity, midgut microbiota, and malaria parasites.

## Results

### Establishment of *REL2* mutants

To explore the role of REL2 in physiological mosquito immune responses, we used CRISPR-Cas9 technology for targeted mutation of the *REL2* locus (*REL2* ^-/-^) by inserting a *3xP3::GFP::SV40* cassette into the sequence encoding the Rel Homology Domain (RHD) (Fig. 1A, B).

**Figure 1:**
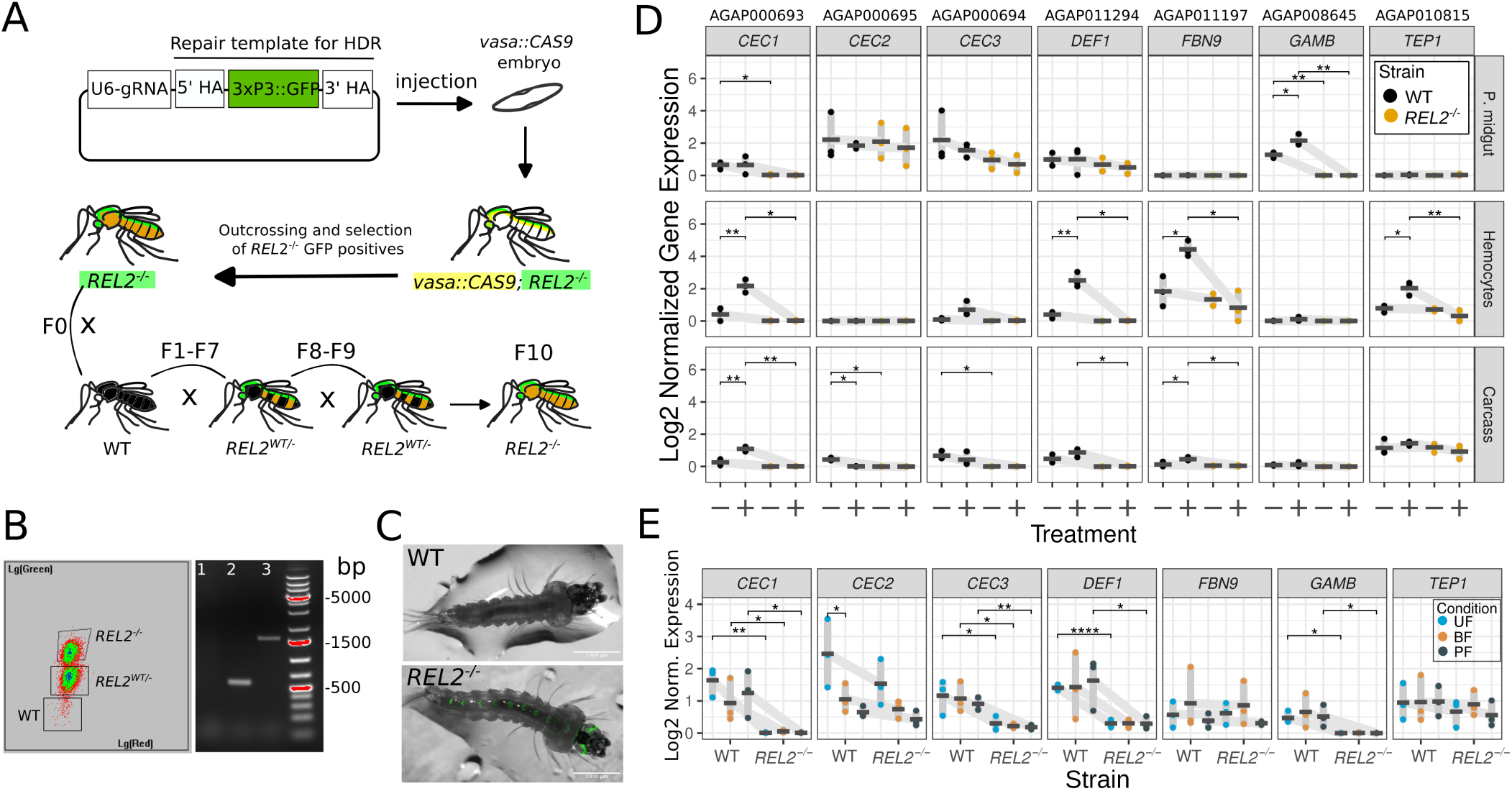
Establishment and phenotyping of *REL2* ^-/-^ mutant line. (A) Schematic summary of the mutagenesis process. A plasmid carrying a guide RNA expressing module (U6-gRNA), together with a repair template for homologous directed repair (HDR) was injected into the embryos of transgenic mosquitoes expressing *vasa::Cas9* ^48^. The generated *REL2* ^-/-^ mutant was outcrossed with WT mosquitoes over 7 generations (F_0_-F_7_). Homozygotes were established by crossing heterozygote *REL2* ^-/-^ individuals for three generations (F_8_-F_10_). (B) Selection of *REL2* ^-/-^ mutants by fluorescence using Large Object Flow Cytometry (left panel) and by PCR genotyping (right panel). Lanes: 1-water control, 2-WT, 3-*REL2* ^-/-^. (C) Microscopy of GFP-negative WT and GFP-positive *REL2* ^-/-^ larvae (note the green fluorescence in the larval eyes). (D, E) RT-qPCR quantification of immune gene expression in the WT and *REL2* ^-/-^ mosquitoes, normalized to the *RPS7* gene. Dots represent independent experiments (N=3, n=1 (pool of 10 mosquitoes)). Crossbars represent mean expression levels, and lines connect the means of expression in the WT and mutant mosquitoes, grouped by experimental condition. Statistical significance was evaluated using Student’s t-test; only significant values are shown: *, *p<*0.05; **, *p<*0.01. (D) Mosquitoes were injected with PBS (-) or PBS with a bacteria mix (*E. coli* and *S. aureus*) (+), four hours post-injection, tissues were dissected: posterior midgut (top row), hemocytes (middle row), and abdominal carcass with fat body (bottom row). (E) Mosquitoes were collected before blood feeding (unfed, UF) and 24 hours post-blood-feeding (BF) and *P. falciparum* infection (PF).

The impact of *REL2* knockout (KO) on the expression levels of potential REL2 target genes was assessed following control thoracic injection of phosphate-buffered saline (PBS) or a bacterial mixture of *Escherichia coli* and *Staphylococcus aureus*. Hemocytes, posterior midgut, and abdominal carcass (containing the fat body) were collected four hours post-injection, and the expression of a set of known immune genes was quantified using RT-qPCR.

Bacterial injection significantly increased the transcript levels of *CEC1*, *DEF1*, *FBN9* and *TEP1* in hemocytes, while *CEC1* and *FBN9* were up-regulated in the carcass/fat body. Notably, compared to control, bacterial injection significantly elevated *GAMB* transcript levels in the posterior midgut but not in the other tissues tested (Fig. 1D). These transcriptional changes induced by bacterial injection were absent in *REL2* mutants, confirming that CRISPR/Cas9-mediated genetic modification resulted in a loss-of-function phenotype. Interestingly, bacterial injection decreased the expression levels of *CEC2* in the carcass of wild-type mosquitoes but this effect was not observed in *REL2* mutants.

Overall, our observations revealed that the tested genes displayed unique tissue-specific expression patterns. Specifically, *CEC1*, *DEF1*, and *CEC3* transcripts were detected in all three analyzed tissues, *TEP1* was found in hemocytes and the carcass, *GAMB* and *CEC2* were primarily observed in the posterior midgut, while *FBN9* was detected predominantly in hemocytes (Fig. 1D).

We then evaluated the impact of *REL2* KO on the expression of the selected genes following blood feeding (BF) and *P. falciparum* infection. Extracts from whole wild-type or *REL2* ^-/-^ mosquitoes were collected before (unfed, UF) and 24 hours after either BF or *P. falciparum* (PF) infection. While BF and *Plasmodium* infection did not significantly affect gene expression, *REL2* KO decreased the expression levels of *CEC1*, *GAMB*, *DEF1*, *CEC3* in all conditions. A modest reduction in *CEC2* expression was observed in unfed *REL2* ^-/-^ compared to wild-type mosquitoes (Fig. 1E). These results confirmed the loss-of-function phenotype of the *REL2* mutants. Considering the tissue-specific expression of the genes tested, further transcriptional analyses were performed using tissue-specific RNA sequencing to obtain a broader picture of the role of the REL2 pathway in mosquito responses to blood feeding and *P. falciparum* infection.

### Identification of the REL2-regulated tissue-specific processes by transcriptome sequencing

To understand the role of REL2 in mosquito responses to blood feeding and *P. falciparum* infection, we examined wild-type (WT) and *REL2* ^-/-^ mosquitoes under different conditions: before blood feeding (unfed, UF) and 24 hours after feeding with either uninfected blood (BF) or blood infected with *P. falciparum* (PF). Our analysis focused on four key immune tissues: hemocytes (hemo), fat body, the anterior midgut with cardia, and the posterior midgut, which is the site of *Plasmodium* ookinete invasion (Fig. 2A).

**Figure 2:**
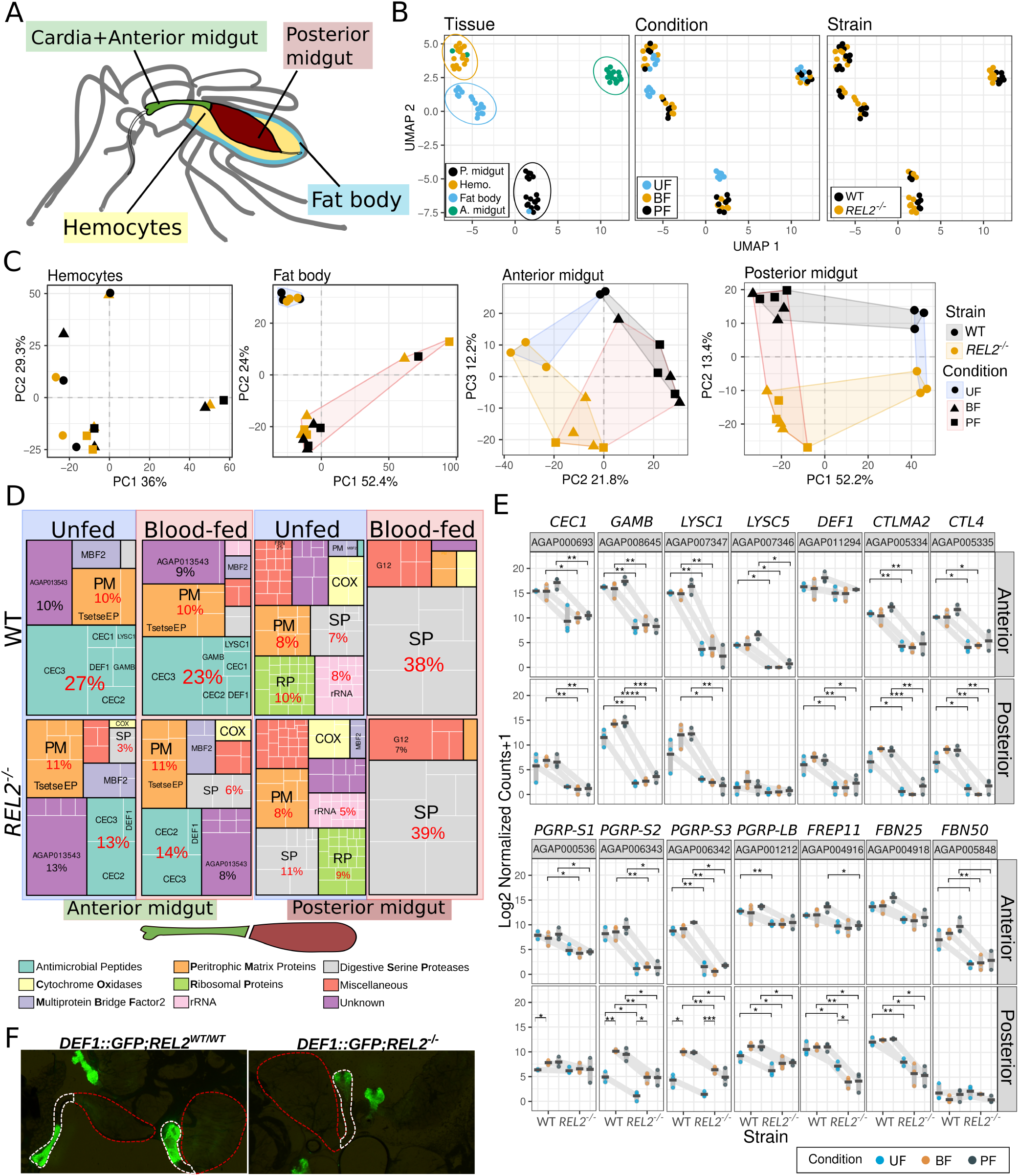
The REL2 pathway regulates immune gene expression in *A. gambiae* midgut. (A) Schematic representation of mosquito tissues used in transcriptomic analyses. (B) UMAP plots of RNA-seq data with identified clusters (circled) colored by tissue (first panel), feeding condition (middle panel) and mosquito strain (third panel). Each dot represents one sample. (C) PCA results of the RNA-seq data, performed separately on each tissue using the 500 most variable genes. Samples are grouping by strain and feeding condition. (D) Treemaps depicting the top 50% of the transcriptome (in transcripts per million, TPM), grouped and colored by functional category. Each square with a white outline represents the expression proportion of a single gene. (E) Immune genes down-regulated in the *REL2* ^-/-^ anterior and/or posterior midgut, identified by *DESeq2*, *P_adj_<*0.05. Statistical significance was evaluated using the Student’s t-test (*, *p<*0.05; **, *p<*0.01; ***, *p<*0.001; ****, *p<*0.0001). UF: unfed; BF: blood-fed; PF: *P. falciparum* infected. Dots represent independent experiments. Mean expression levels (crossbars) for each mosquito line are grouped by experimental conditions and connected by gray lines. (F) Fluorescence microscopy images of dissected midguts from WT (*DEF1-GFP*;*REL2^W^ ^T/W^ ^T^*, left panel), and *REL2* mutant (*DEF1-GFP*;*REL2* ^-/-^, right panel) genetic backgrounds illustrating the expression of the *DEF1::GFP* reporter gene. Dashed lines outline the cardia and anterior midgut (white) and the posterior midgut (red).

Uniform Manifold Approximation and Projection (UMAP) analysis of the entire dataset identified four distinct clusters of mosquito samples, each corresponding to a specific tissue type (Fig. 2B, left panel). However, a few mismatched samples were observed: two anterior midgut samples grouped with hemocytes, and one fat body sample clustered with the posterior midgut group. Additionally, transcriptomes of unfed and blood fed samples (infected or not) showed clear separation for the fat body and posterior midgut samples (Fig. 2B, middle). Notably, differential clustering between WT and *REL2* ^-/-^ strains was detected only for the anterior and posterior midgut samples (Fig. 2B, right panel), highlighting tissue-specific effects of REL2 deficiency.

To confirm tissue-specific clustering, we selected 20 highly expressed genes in WT mosquitoes (UMAP, *P_adj_<*0.05), and plotted their expression patterns for each tissue (Fig. S1A). Expression of most of these genes matched the previously reported tissue-specific patterns, validating the specificity of our transcriptome analyses. The hemocyte transcriptome was enriched with genes encoding prophenoloxidases (PPOs) and CLIP-serine proteases, which are involved in melanization processes, and with fibrinogen-like proteins (FBN)^39^. The fat body transcriptome was primarily characterized by genes encoding Leucine-Rich Repeat proteins and those involved in lipid metabolism. Consistent with previous reports, the transcriptome of the cardia and anterior midgut was dominated by genes encoding AMPs, the putative transcriptional activator Multiprotein Bridge Factor 2 (MBF2), the peritrophic matrix protein Tsetse EP and carbohydrate digestive enzymes (Fig. S1A, B)^27^. In the posterior midgut, genes coding for inducible digestive trypsins and carboxypeptidases were highly enriched, particularly after blood feeding (Fig. S1A, C). As previously reported, expression levels of serine protease genes in the unfed *A. gambiae* posterior midgut were approximately 2.5-fold lower than those in *Ae. aegypti* ^27^. However, this difference was fully compensated after blood feeding, with a significant increase in serine protease expression reaching levels comparable to those observed in *Aedes*.

Next, we used Principal Component Analysis (PCA) to investigate the effect of blood feeding on tissue-specific gene expression (Fig. 2C). No differences were observed in hemocyte samples under any condition. However, a clear separation was detected between unfed and blood-fed (infected or not) samples in the fat body, anterior and posterior midgut (Fig. 2C). Additionally, PCA revealed significant transcriptional differences between WT and *REL2* ^-/-^ mosquitoes in the anterior and posterior midgut. In contrast, no notable differences were detected in fat body and hemocytes. These results suggested that REL2-mediated regulation is more pronounced in midgut-associated immune responses.

To connect the observed transcriptional changes with potential biological functions, we performed GO-term analyses on differentially-expressed genes (DEGs) detected after blood feeding in WT mosquitoes. In the fat body and posterior midgut of unfed mosquitoes, GO analyses revealed enrichment in such processes as RNA processing, ribosome biogenesis, and related metabolic and catalytic activities (Fig. S1D). In particular, protein and lipid transport, and glycosylation were enriched in the fat body; fatty acid beta-oxidation, iron ion homeostasis, and intracellular cholesterol transport in the posterior midgut; and lipid transport, proteolysis, and peptidase activity in the extracellular space were enriched in the anterior midgut (Fig. S1D). The fat body showed the strongest response to blood feeding, with 492 up-regulated genes. In the posterior midgut we identified 231 up-regulated genes, while 136 genes were up-regulated in the anterior midgut. The processes that were primarily induced by blood feeding in these tissues were related to metabolism and digestion rather than immunity.

To investigate the role of REL2 in biological processes within midgut regions, we compared transcriptional activity between WT and *REL2* ^-/-^ mosquitoes. We adapted a data visualization technique from a recent study ^27^, which represents transcript read counts per million (TPM) as a treemap. This approach displays the top 50% of the transcriptome for each tissue, mosquito strain and feeding condition (unfed or blood-fed), categorized by function. In WT mosquitoes, notable differences in transcriptional complexity were observed between the anterior midgut (including the cardia) and the posterior midgut. In the anterior midgut, just 10 genes accounted for the 50% of total transcriptome. Among these, 27% were AMP genes, 10% encoded peritrophic matrix proteins, 10% corresponded to the unknown gene AGAP013543, and 3% were attributed to *MBF2*, a potential transcriptional factor (Fig. 2D)^27^. *REL2* KO led to a two-fold reduction in AMP gene expression (Fig. 2D left panel), while expression levels of other functional categories, such as peritrophic matrix proteins, remained unchanged. Although *REL2* deficiency did not significantly alter transcriptional investment after blood feeding, it resulted in a two-fold increase in the expression of genes encoding serine proteases (SP) at the expense of AGAP013543 (Fig. 2D).

In contrast to the highly specialized expression profile of the anterior midgut, 50% of the posterior midgut transcriptome included 126 transcripts encoding such protein families as ribosomal proteins, digestive serine proteases, cytochrome oxidases (COX) and peritrophic matrix proteins. Blood feeding induced a significant shift in the transcriptional repertoire, favoring enrichment of transcripts encoding digestive serine proteases (SP, 38%) and the protein of unknown function G12 (4%) (Fig. 2D). This shift was accompanied by a reduction in the expressional investment into ribosomal proteins, rRNA, peritrophic matrix proteins and COX. The impact of blood feeding on the transcription of digestive enzyme genes in the posterior midgut was evident across the entire transcriptome (Fig. S1C). Similar changes were observed in *REL2* mutants, with the exception of G12, whose expression was two-fold higher than in WT mosquitoes. Notably, the relatively modest investment in AMPs observed in unfed WT mosquitoes was lost following the blood feeding and in *REL2* mutants. These findings suggest that REL2 plays a crucial role in regulating AMP gene expression in both compartments of the midgut (Fig. 2D).

We next examined REL2-mediated processes in the midgut tissues at the individual gene level. Out of 75 DEGs identified in the anterior midgut and cardia of *REL2* mutants, 60 were downregulated and 15 up-regulated. Posterior midgut in *REL2* mutants featured 133 DEGs (81 down-regulated and 52 up-regulated). Gene Ontology (GO) analysis revealed that REL2 deficiency resulted in significant suppression of immune- and antibacterial-related processes in both midgut sections (Fig. S2A). We also observed changes in other GO categories, such as metabolic processes in the posterior midgut and detoxification processes in the anterior midgut. However, these transcriptional changes following the blood meal were rather modest, particularly involving detoxification processes.

The GO category most profoundly affected by *REL2* deficiency was the expression of AMP-encoding genes, including *GAMB*, *CEC1*, two C-type lysozymes (*LYSC1* and *LYSC5*), and *DEF1* (Fig. 2E, S3A). This category was accompanied by altered expression levels of genes encoding C-type lectins (CTLMA2 and CTL4), PGRP receptors (short PGRP-S1, -S2, -S3, and long PGRP-LB), fibrinogen-like proteins (FREP11 and FBN25), and the serine-protease inhibitor 9 (SRPN9). While the anterior midgut with cardia exhibited the highest AMP expression levels among all sequenced tissues (Fig. S1A,B, S3A), the posterior midgut also showed strong immune gene expression, including *GAMB*, *LYSC1*, *CTLMA2*, *CTL4*, *PGRP-S2*, -*S3*, *FREP11* and *FBN25* (Fig. 2E, S1B).

Our analysis highlighted high levels of constitutive expression of immune genes in the anterior midgut with cardia, regardless of feeding condition. In the posterior midgut, blood feeding strongly induced expression of *PGRP-S2*,-*S3*, *LRIM4* and *TEP3*, while the expression of *GAMB*, *LYSC1*, *PGRP-LB*, *CTLMA2* and *CTL4* was mildly up-regulated by blood feeding (Fig. 2E, S2B). Moreover, expression of *FBN25* and *FREP11* was down-regulated following blood feeding in WT mosquitoes, while *REL2* deficiency further decreased expression levels of these fibrinogens. Interestingly, expression of *LYSC5*, *PGRP-S1*, *FBN50* (Fig. 2E), *CEC3*, *SRPN9* and *FREP12* (Fig. S2B) was regulated by REL2 only in the anterior midgut and cardia. Conversely, REL2-mediated induction of *LRIM4* and *TEP3* after blood feeding was only detected in the posterior midgut (Fig. S2B).

Unexpectedly, the constitutive expression of *PGRP-LC*, encoding the transmembrane receptor of the REL2 pathway, in the anterior midgut and cardia - as well as blood-induced upregulation in the posterior midgut - was independent of REL2 (Fig. S3B). These results suggested that *PGRP-LC*, along with other intracellular pathway components, is not regulated by the pathway itself. Interestingly, we observed higher levels of *REL2* transcripts in *REL2* ^-/-^ mutants (Fig. S3B), however most of the sequencing reads were mapped to the *REL2* transcript region upstream of the gRNA target site (Fig. S3C), and corresponded to truncated non-functional *REL2* transcripts.

Previous reports described constitutive expression of AMP genes in the cardia^26,27,40^. To visualize *in vivo* the expression of the REL2-target gene *DEF1* in the midgut, we utilized a mosquito transgenic reporter line that expresses *GFP* under the *DEF1* promoter (*DEF1::GFP)* ^41^. We observed the strongest GFP signal in the cardia and adjacent half of the anterior midgut, with only a low, diffuse signal in the distal anterior midgut region and posterior midgut (Fig. 2F, first panel). To further investigate the role of REL2 in regulating *DEF1* expression, we crossed this reporter line with a *REL2* deficient line *REL2* ^-/-^ to create a homozygous *DEF1::GFP;REL2^-/-^* line. In contrast to the GFP expression observed in WT mosquito midguts, the *DEF1::GFP;REL2^-/-^* mosquito midguts showed GFP expression only in the cardia. The anterior midgut lost reporter fluorescence in REL2 absence (Fig. 2F, second panel). These results support the transcriptional data and demonstrate that REL2 regulates *DEF1* expression in the anterior midgut but that the constitutive expression of *DEF1* in cardia is REL2-independent.

Finally, we examined the transcriptional responses mediated by REL2 at 24 h after *P. falciparum* infection. Only eight genes were differentially expressed in the posterior midgut of *REL2* mutants compared to WT (Fig. S2C). Among these, two genes were significantly down-regulated in REL2 mutants after blood feeding and *P. falciparum* infection: *Mucin-2* ^42^ and a gene encoding a peritrophic matrix-associated protein^43^. Notably, *P. falciparum* infection but not blood feeding, modestly but significantly induced the expression of these genes in *REL2* mutants. Additionally, three genes - *syntaxin 16*, *carbonic anhydrase* and *scavenger receptor class A* - were modestly down regulated by parasite infection in *REL2* mutants. Conversely, the expression of *Rab-8A*, *Rhomboid4B* and *Nucleolar Complex protein3* was modestly upregulated. These results suggested that although *P. falciparum* midgut colonization did not trigger strong REL2-mediated transcriptional responses at 24 hours after infection, they did highlight the role of REL2 in regulating certain components of the peritrophic matrix, vesicular transport, and proteins putatively involved in tissue stress response.

We concluded that the observed REL2-mediated transcriptional regulation of upstream sensing module and the downstream effector AMPs in the midgut before and after blood feeding provides a powerful model to investigate the functional consequences of REL2 depletion on mosquito survival and susceptibility to *P. falciparum*.

### *REL2* depletion promotes midgut bacteria growth and mosquito lethality without affecting susceptibility to *P. falciparum*

WT mosquitoes and *REL2* mutants were infected with *P. falciparum* and 11 days later the percentage of infected mosquitoes (infection prevalence) and the number of oocysts (infection load) were quantified in the dissected midguts. High variability was observed between experimental repeats and no significant difference between WT and *REL2* mutants was detected (Fig. 3A). However, we noted the increased mortality rates in the *REL2*-deficient group. To test whether the death of immuno-compromised mosquitoes was induced by *P. falciparum* infection, we quantified female survival after blood feeding or *P. falciparum* infection. We found that the increased lethality in *REL2* mutants was due to blood feeding and not *P. falciparum* infection (Fig. 3B), regardless of the infection outcome (Fig. 3C).

**Figure 3:**
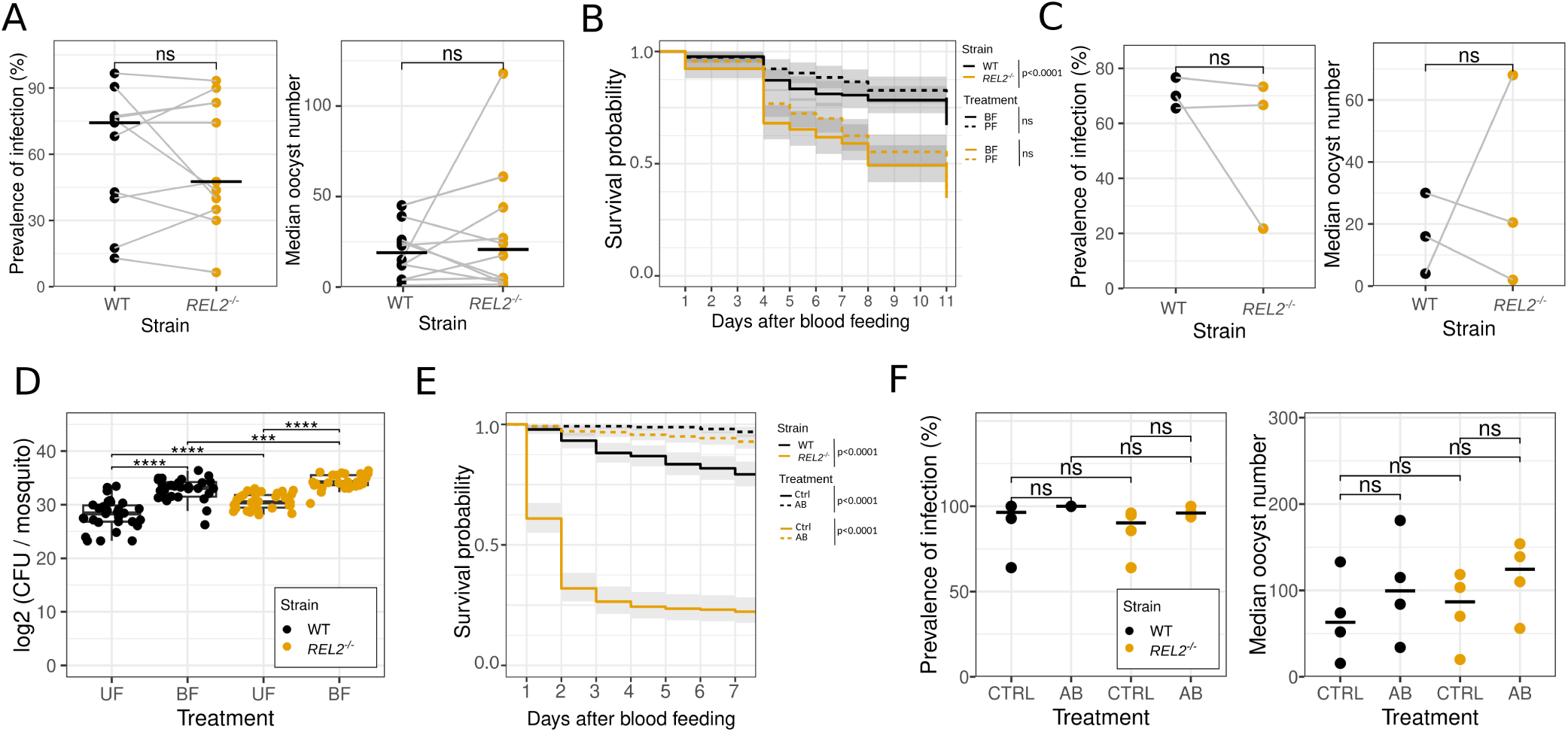
Immune suppression causes bacterial overgrowth and increased mortality in *REL2* ^-/-^ mosquitoes, without affecting susceptibility to *P. falciparum*. (A, C, F) Prevalence of *P. falciparum* infection (%) and median oocyst count per experiment for WT (black) and *REL2* ^-/-^ (yellow) mosquitoes infected with at least one oocyst. (A) Experimental repeats are shown by lines, crossbars indicate the median of infection rates (N=10, n=18-40). (B, E) Kaplan-Meier survival analysis of WT and *REL2* ^-/-^ mosquitoes: (B) post-blood-feeding and *P. falciparum* infection (N=3, n=30-70); (E) post-blood-feeding with (AB) or without (CTRL) antibiotic treatment (N=3, n≥50). (C) Results of *P. falciparum* infection in female mosquitoes as shown in the survival experiments in (B) (N=3, n=12-30). (D) Bacterial loads expressed as colony forming units (CFUs) from plated mosquito homogenates, isolated from unfed mosquitoes or from mosquitoes 24 h after blood feeding. Dots represent CFU counts for individual mosquitoes (N=3, n=10). (F) Prevalence of *P. falciparum* infection and oocyst loads in untreated (CTRL) and antibiotic-treated (AB) WT and *REL2* ^-/-^ mosquitoes (N=4, n=15-30). (A, C, D, F) Statistically significant differences were evaluated using the Wilcoxon Rank Sum test, p*>*0.05, ns; ***, *p<*0.001; ****, *p<*0.0001. (B, E) Statistically significant differences were evaluated with Log-rank test.

The high variability in infection outcomes and the lethality of mutant mosquitoes induced by blood feeding suggested that both phenotypes may result from uncontrolled bacterial proliferation in the midgut. Therefore, we compared bacterial loads in WT and *REL2* mutants by counting colony-forming units (CFUs) from plated homogenates of whole mosquitoes collected before (unfed) and 24 h after blood feeding (BF). We found significantly higher bacterial load in the *REL2* mutants compared to WT mosquitoes, both before and after blood feeding (Fig. 3D), indicating that immunosuppression caused bacterial overgrowth. To confirm that mortality observed in *REL2* mutants was caused by bacteria, we treated WT and *REL2*-deficient females with sugar meals containing a mix of antibiotics before blood feeding. The antibiotic treatment fully rescued mutants’ survival and also improved the survival of WT mosquitoes, particularly in experiments with low survival rates in both groups (Fig. 3E).

We re-examined the role of REL2 depletion on mosquito susceptibility to *P. falciparum* infection after antibiotic treatment. Both WT and *REL2* ^-/-^ mosquitoes were administered antibiotics before infection. We then quantified the prevalence and intensity of infection by measuring oocyst loads in the dissected mosquito midguts, as described above. Antibiotic treatment reduced interexperimental variability and slightly increased both infection prevalence and oocyst loads. Importantly, under these conditions, no statistically significant differences were detected between WT and *REL2* mutants (Fig. 3F).

Overall, our findings suggested that REL2-mediated immune responses in the midgut protect mosquitoes from lethal bacterial infections, likely through antimicrobial peptide activity. However, we found no supporting evidence for REL2 function in limiting *P. falciparum* development in the midgut. Instead, our results demonstrate the contribution of midgut microbiota to the interexperimental variability often observed in the outcomes of *P. falciparum* infections in mosquitoes.

### *Serratia* spp.-induced dysbiosis hinders survival and *P. falciparum* development in *REL2* deficient mosquitoes

Depletion of REL2 leads to excessive bacterial growth and mosquito mortality. To identify the bacterial species responsible for these effects, we sequenced PCR amplicons of the variable region 4 (V4) of the *16S* rDNA gene, using DNA isolated from the midguts of WT and *REL2* mutant mosquitoes at 24 h after blood feeding. To correlate microbial composition with survival rates, survival rates of the remaining mosquitoes from both experimental groups were also evaluated.

In five independent experiments, we observed a wide range of mosquito survival rates ranging from very poor to very good (Fig. 4A, left panel). A detailed analysis of bacterial communities from these experiments revealed a strong correlation between low survival rates and high prevalence and abundance of *Serratia spp.* (Fig. 4A, right panel, experiments 1 and 2). In contrast, mosquitoes from experiments with high survival rates (experiments 3, 4, albeit not experiment 5) showed more diverse microbial communities.

**Figure 4:**
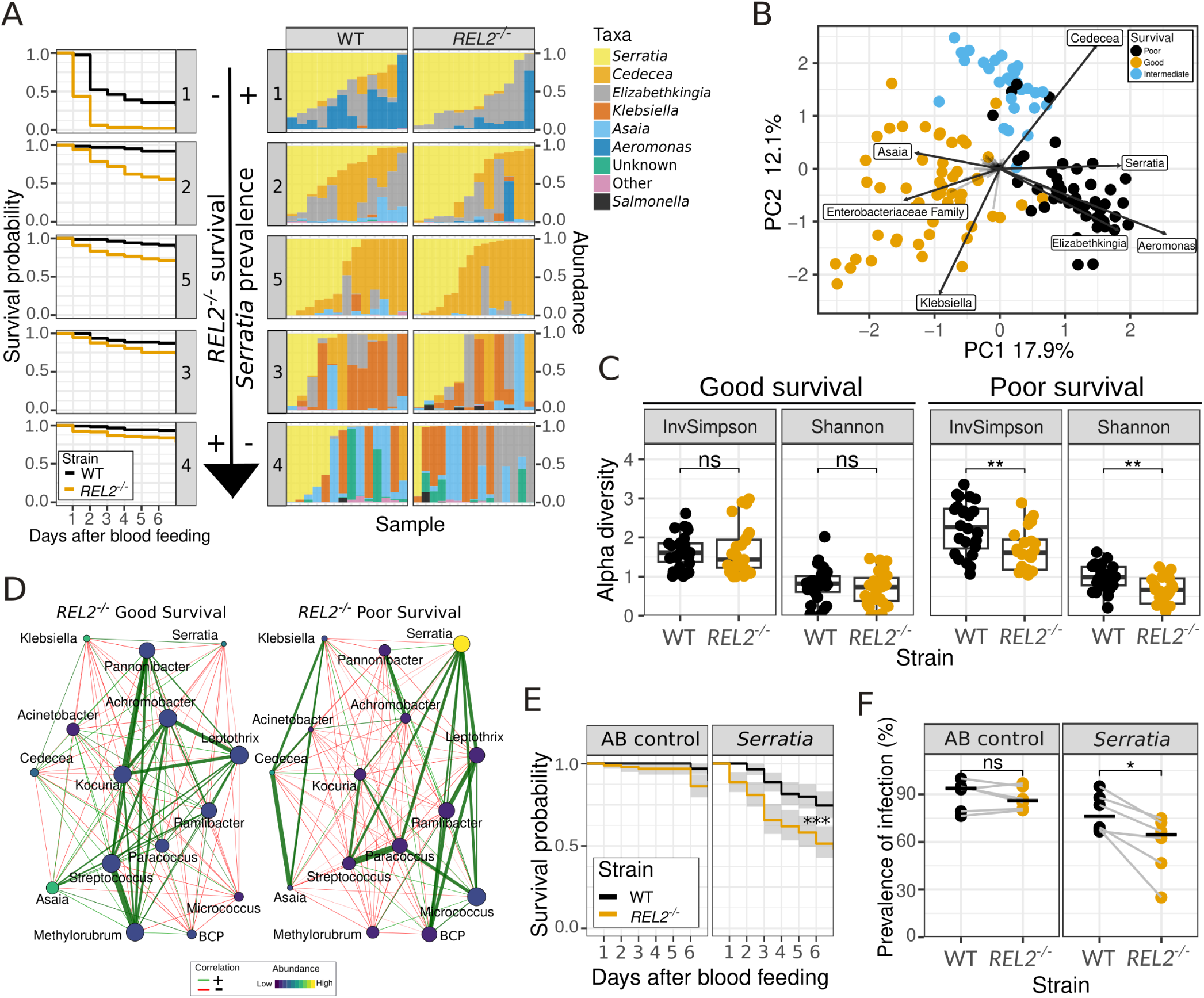
Impact of *Serratia* spp. on *Anopheles* survival and *P. falciparum* infection in *REL2* mutants. (A) The survival of *REL2* ^-/-^ mutants (left panel, N=5, n≥60), is correlated with the prevalence and relative abundance of *Serratia* in the mosquito midguts (right panel, (N=5, n∼15), as determined by 16S rDNA V4 locus sequencing 24 h post-blood-feeding. Bar plots display the microbiome composition (aggregated at the genus level) from individual mosquitoes. (B) PCA plot illustrating the *β*-diversity of the mosquito microbiome. Samples are color-coded based on survival phenotype, with arrows indicating the most important taxa (loadings) for each survival phenotype. (C) *α*-diversity plots from experiments with high survival rates (Experiment 3 and 4) and low survival rates (Experiments 1 and 2). (D) Comparison of microbial association networks estimated using Pearson correlation coefficients (*>*0.3) for *REL2* ^-/-^ mutants from experiments with high and low survival rates. The size of each node represents the eigenvector, and color indicates relative abundance according to the heatmap. Positive correlations between the taxa are shown by green lines, red lines indicate negative correlations. The layout combines the two networks. (E, F) The effect of *Serratia* sp. Ag2 colonization on mosquito survival and *P. falciparum* infection. Midguts of WT and *REL2* ^-/-^ mosquitoes were treated with antibiotics, and then orally colonized with *Serratia* sp. Ag2 (OD_600_=0.5) in a sugar meal (*Serratia* group) or left uncolonized (AB control). (E) Kaplan-Meier survival analysis comparing *Serratia*-colonized mosquitoes to AB control over 7 days post-blood-feeding (N=3, n≥20). Statistically significant differences were assessed by the Log-rank test, ***, *p<*0.0001. (F) Prevalence of *P. falciparum* infection (oocysts, %) in the midguts of *Serratia*-colonized mosquitoes *versus* AB control. Horizontal bars show medians (N=6, n=12-50). (C, F) Statistically significant differences were evaluated with the Wilcoxon Rank Sum test (ns, *p>*0.05; *, *p<*0.05; **, *p<*0.01).

Principal Component Analysis (PCA) was used to compare bacterial communities across multiple dimensions based on *β*-diversity. This analysis provided quantitative support to the hypothesis that samples cluster according to survival phenotype (Fig. 4B). Poor mosquito survival correlated with the presence of *Serratia* spp., *Aeromonas* spp. and *Elizabethkingia* spp. (Fig. 4B). Conversely, good mosquito survival was linked to the presence of *Asaia* spp., *Klebsiella* spp. and a species from the Enterobacteriaceae family. Overall, the *β*-diversity of the midgut bacteria varied between independent experiments but showed no difference between WT mosquitoes and *REL2* mutants within the same experiment (Fig. S4A).

The *α*-diversity indices measure species diversity within a single sample and are often used to assess the overall health and stability of the microbiome. Significant differences in the *α*-diversity index were observed in *REL2* mutants when comparing mosquitoes with low and high survival phenotypes (Fig. 4C). These results suggest a correlation between low survival rates of the *REL2* mutants and reduced diversity of their midgut microbiomes, dominated by *Serratia* spp., *Aeromonas* spp. and *Elizabethkingia* spp..

Both *Serratia* and *Elizabethkingia* species were identified in the midgut bacterial communities of *REL2* mutants. We examined whether the potential co-occurrence of these bacteria impacted mosquito survival. To do this, we performed a microbial association network analysis using microbial abundance data from the samples of REL2-deficient mosquitoes under conditions of high and low survival. We focused on the 30 most prevalent bacterial genera and calculated Pearson’s correlation coefficients among them. Notably, in bacterial communities found in the samples with low survival, we identified positive associations between the highly abundant *Serratia* and several rare taxa (Fig. 4D). In contrast, the network linked to high survival showed strong associations among rare taxa, while moderately abundant *Serratia* and *Asaia* displayed only weak connections. These results suggested that high *Serratia* spp. abundance was a probable driver of the poor survival of *REL2* mutants, and indicated that both genotype and environment contribute to the survival outcome.

To directly examine the potential impact of *Serratia* spp. on mosquito mortality, we fed both WT mosquitoes and *REL2* mutants with two concentrations (OD_600_= 0.5 and 1) of *Serratia* sp. Ag2 strain which was isolated from *A. gambiae* ^44^. Successful colonization was confirmed by plating mosquito homogenates on agar plates and measuring colony-forming units (CFU). Two days after colonization, no significant differences were detected in bacterial loads between the two mosquito strains (data not shown). While non-colonized control mosquitoes from both strains showed high survival rates, those colonized with *Serratia* sp. Ag2 experienced significantly higher mortality, particularly in REL2 mutants, at both bacterial doses (Fig. 4E, S4B). These results directly support the observed correlation between the presence of *Serratia* spp. in the midgut and high mortality in *REL2* mutants.

Previous field and laboratory studies have documented the impact of *Serratia* colonization on *Plasmodium* infection^19,45–47^. In this study, we examined how *Serratia* sp. Ag2 strain affects the susceptibility of WT and *REL2* mutant mosquitoes to *P. falciparum* infection. Mosquitoes were colonized with *Serratia* sp. Ag2 (OD_600_= 0.5) as described above and infected with *P. falciparum*. While similar infection prevalence was observed in both WT and *REL2*-deficient mosquitoes treated with antibiotics, colonization with *Serratia* significantly reduced *P. falciparum* prevalence in *REL2* mutants compared to WT mosquitoes (Fig. 4F). However, no significant differences were detected on oocyst loads (Fig. S4C). These findings suggest that the immunosuppression resulting from REL2 depletion may indirectly impact mosquito susceptibility to *P. falciparum* infection, and that *Serratia* sp. Ag2 negatively impacts parasite development within the mosquito midgut.

In conclusion, our results identified the critical role of REL2 in maintaining midgut microbial homeostasis after blood feeding. This function of REL2 is vital for restricting overgrowth of opportunistic pathogens, such as *Serratia* spp., which can also impede *P. falciparum* development in the mosquito midgut.

## Discussion

The immune system is essential for protecting organisms from daily threats posed by invading microbes. Studies in insects have traditionally focused on immune responses to experimental infections with model pathogens, often using artificial feeding or injection methods. Here, we explored the physiological role of the immune system after natural microbial expansion induced by blood feeding in both wild-type and REL2-deficient mosquitoes. Our findings revealed a critical role of REL2 in maintaining midgut microbial homeostasis. Disruption of REL2 signaling led to a shift in bacterial communities, favoring *Serratia* spp., and increased mosquito mortality, highlighting the importance of effective immune signaling and the impact of environmental bacterial opportunists on mosquito physiology. In contrast to earlier studies^30,34^, but in line with more recent research ^33^, our results do not support a direct role of REL2-mediated immune responses against *P. falciparum* parasites. Instead, they suggest that the midgut abundance of *Serratia* spp. directly impairs the efficiency of *Plasmodium* colonization in mosquito midguts.

Fat body expressed AMP genes are the major targets of the *Drosophila* IMD pathway after a bacterial injection. Reported here transcriptional analyses of the cardia, anterior and posterior midguts of *Anopheles* mosquitoes showed that blood feeding induced REL2-mediated expression of AMPs only in the posterior midgut. In contrast, the cardia and the proximal part of the anterior midgut exhibited strong, constitutive expression of AMPs that was independent of REL2. While our results suggested that REL2-dependent regulation of AMP expression was crucial for maintaining bacterial homeostasis after a blood meal, further research is needed to determine specific roles of individual AMPs in counteracting dysbiosis caused by *Serratia* spp. overgrowth.

Until recently, the antibacterial properties of AMPs were believed to rely on a broad additive or synergistic mode of action. While some AMPs may indeed contribute to this broad activity ^49,50^, two specific *Drosophila* AMPs have been reported to exhibit antibacterial activity against particular bacterial species. For instance, Diptericin has been shown to be crucial in defenses against *Providencia rettgeri*, but not other *Providencia* species^49^. Conversely, Attacin protected the fruit flies against *Serratia marcescens* ^51^. Notably, our data showed that the expression of *Anopheles Attacin* is not regulated by REL2. Consequently, this AMP is dispensable for protection against *Serratia* spp., while other REL2-targeted AMPs mediate *Anopheles* resistance to *Serratia* overgrowth in the posterior midgut.

Our findings revealed significantly higher AMP expression levels in the cardia and anterior midgut of *Anopheles* as compared to *Aedes* mosquitoes^52^. It remains to be explored whether this difference reflects an evolutionary adaptation of the mosquito species to opportunistic pathogens. Another intriguing observation from this study is the potential involvement of REL2-regulated fibrinogen-like (FBN) proteins in midgut antibacterial responses. Although the expression of these FBNs has been reported to increase in response to bacterial infections, they have not yet been functionally characterized^48,53,54^.

Interestingly, we did not observe REL2-mediated positive feedback loop for the intracellular components of this signaling pathway starting from the pathway receptor PGRP-LC to the negative regulator Caspar. Instead, we identified REL2-mediated regulation of potential upstream components, such as PGRPs and CTLs. These modulators of immune signaling can be classified as either positive or negative regulators. Positive regulators bind to microbial PGNs and activate the pathway, while negative regulators catalyze or compete for PGNs to balance immune activation. PGRP-LB is one of the known negative regulators of REL2 signaling in the midgut ^37^. Additionally, the short PGRPS, PGRP-S2 and PGRP-S3, are predicted to have amidase activity, potentially hydrolyzing PGNs to attenuate the quantities of activating ligands. The specific expression of these short PGRPs in the midgut aligns with earlier reports suggesting that they are not essential for mosquito defenses against systemic bacterial infections in the hemolymph^34^. Notably, RNAi-mediated knockdown of these genes moderately increased the prevalence of *P. falciparum* infections ^37^. In light of our findings, it is likely that the observed increase in *P. falciparum* prevalence may be due to an indirect effect on parasite development through dysregulation of REL2 pathway and bacterial dysbiosis. Unlike PGRP-S2 and -S3, PGRP-S1 lacks a conserved amidase signature and may activate the REL2 pathway by binding to peptidoglycan^55^. Further studies are needed to confirm this hypothesis.

We also detected REL2-mediated expression of *CTL4* and *CTLMA2* in the anterior and posterior midgut. These lectins form a dimer in the hemolymph^56^, which negatively regulates the melanization of *Plasmodium* parasites and fungi^57,58^. Additionally, CTL4/CTLMA2 dimer protects *A. gambiae* against injected Gram-negative bacteria by an unknown melanization-independent mechanism^58,59^. These two functions occur in the hemolymph and are REL2-independent. Our data indicate that CTL4/CTLMA2 complex may potentially function as a negative regulator of REL2 signaling in the midgut, as *CTL4* knockdown was reported to decrease mosquito bacterial loads after blood feeding^58^. However, the molecular mechanisms underlying CTL4/CTLMA2-mediated regulation remain unknown.

Due to the tissue-specific regulation of AMPs in the cardia, anterior, and posterior midgut, we propose a two-step model for the role of REL2 in limiting bacteria proliferation. High AMP expression in the anterior midgut likely functions as a pathogen filter, reducing bacterial entry into the midgut. In the absence of blood feeding, bacterial loads in the posterior midgut remain low. However, during blood feeding, bacterial proliferation produces large quantities of PGNs, which activate the REL2 pathway and trigger AMP synthesis. Increased AMP activity inhibits bacteria growth. A positive feedback loop within the pathway, which controls the expression of negative upstream regulators, terminates the signaling to restore tissue homeostasis. This model challenges the notion of immune tolerance in the posterior midgut ^52^. Indeed, the REL2 pathway is crucial for mosquito survival after blood feeding, despite constitutive AMP expression in the cardia. We propose that REL2 activity in both sections of the midgut is essential for maintaining microbial homeostasis. Consistent with previous reports^15,60–62^, we also observed REL2-dependent responses in hemocytes and the fat body after injection of a bacterial mix. However, at the population level, the role of such sepsis-like wounding in individual mosquitoes appears to be less live-threatening compared to the bacterial proliferation that follows blood feeding.

In our experiments, we did not control bacterial exposure, which mimicked inert-experimental variability of environmental microbiome variability and was essential to unravel the role of REL2 in controlling opportunistic pathogens such as *Serratia* spp.. The role of *Serratia* in driving dysbiosis appears to be conserved across the dipteran order^63^. Some strains of *S. marcescens* have been reported to damage the insect intestinal epithelium^64,65^ and cross into the hemolymph after ingestion^60,64^. In *Anopheles*, the inhibition of *P. berghei* infection by *Serratia* spp. has been linked to flagellar function^45^ and/or to the induction of host immunity^66^. The relationship between these two phenotypes remains unclear. Soluble factors isolated from *S. marcescens* also inhibited asexual stages of *P. falciparum in vitro* ^67^; however, their identity is unknown. In the reported here experimental *P. falciparum* infections, *Serratia* likely inhibited the fertilization of gametes and/or ookinete development, given that oocyst prevalence positively correlates with the efficiency of ookinete maturation^68^.

Unlike in *Drosophila*, where the IMD pathway regulates many immune genes ^69^, we identified only a few major target genes linking REL2 activity to bacterial detection and effector activation. We demonstrate that the REL2 pathway maintains bacterial homeostasis in the midgut and, thereby, indirectly shapes the local environment encountered by the invading parasites. While our study provides valuable conceptual insights, it is limited by the laboratory-specific environment and a bias of laboratory-enriched bacterial species. Future research should explore a wider array of microbial interactions to further elucidate the role of the REL2 pathway role in regulating both healthy microbial communities and *Plasmodium*-inhibiting bacteria occurring in natural mosquito environment. Understanding the precise mechanisms governing interactions among bacteria, mosquito hosts, and the pathogens they transmit holds great promise to uncovering the molecular mechanisms underlying ecology and malaria transmission by mosquito vectors.

## Methods

### Mosquito stocks

*A. gambiae* G3 strain was used as a wild-type (WT) control and for creation of the *REL2^−/−^* mutant line.

The *REL2* ^-/-^ mutant line was obtained by insertion of a *3xP3::GFP* fluorescence cassette into the coding sequence of the *REL2* gene, using the CRISPR/Cas9 system (Figure 1a). Briefly, a plasmid expressing a guide RNA under the control of AGAP013557 U6 promoter, targeting the 5’-*GCAGCAGCAGTGGTCAGTGT*-3’ sequence in *REL2*, and containing *REL2* 3’ and 5’ UTR regions of homology and a *3xP3::GFP* transgenesis marker cassette as a repair template, was injected into embryos of an *A. gambiae* line expressing *vasa::Cas9* transgene ^48^. A GFP reporter line was derived from a single insertion event. The *REL2* ^-/-^ line was back-crossed with WT mosquitoes for more than 7 generations. *REL2* ^-/-^ mutants were selected by GFP fluorescence using Large Object Flow Cytometry (Complex Object Parametric Analyzer and Sorter, COPAS) and their homozygosity confirmed by genotyping (Figure 1b).

To create transgenic *DEF1::GFP* reporter line, 1,341 bp of the *DEF1* (AGAP011294) 5’-UTR sequence including the endogenous *DEF1* initiator codon (*ATG*) and 353 bp of the *DEF1* 3’-UTR sequence as transcription terminator were cloned upstream and downstream of the *GFP* coding sequence, respectively, within a *piggyBac* vector containing *3xP3::DsRed* and *OpIE2::pac* (puromycin resistance gene) transgenesis selection markers. Mosquito transgenesis in the G3 background was performed as described previously^41^.

### Mosquito rearing

Mosquitoes were reared at 28°C 80% relative humidity (RH), with 12 h day-night cycles. WT and *REL2* ^-/-^ larvae were reared in the same water to allow the sharing of microbial communities, separated by a mesh window, unless stated otherwise. Larvae were fed with tuna-liver food mix, adults were fed with 10% sucrose solution, *ad libitum*. Three- to four-day-old females were used for experiments. Mosquitoes which only received sucrose meals, but not blood-meals, were called unfed (UF) through out the manuscript. Blood feeding of mosquitoes was done with human donor blood using artificial membrane feeder system. Only fully engorged females were kept for experiments. The same reconstituted blood was used for non-infectious (BF) and infectious blood feedings with *P. falciparum* (PF) when experiments were done in parallel.

### Infectious blood-feeding

*P. falciparum* NF54 parasites were cultured at 37℃ with 3% O2 and 4% CO2. Asexual blood stages were maintained in a complete culture media (RPMI 1640 medium with L-glutamine, 25 mM HEPES (ThermoFisher), 10 mM hypoxanthine (c-c-pro GmbH), 20 *µ*g/ml gentamicin (Sigma), supplemented with 10% A+ human serum (Haema) in O+ human erythrocytes (Haema) at 2-5% haematocrit. Media was changed daily and the cultures were diluted every two to three days to maintain the parasitaemia below 3-4%. To obtain gametocytes for the infection, the asexual stages were diluted with fresh red blood cells to 1% parasitaemia and 4% haematocrit in 6 ml complete medium without gentamicin, to avoid antibiotic interference during mosquito feeding. Gametocytes were cultured for 14 days with daily medium changes. The number of mature gametocytes was evaluated using a cell counting chamber (Neubauer). Mature gametocyte cultures (15-16 days post culture set-up) were harvested by centrifugation and mixed with pre-warmed human red blood cells and serum to obtain final gametocytemia of 0.15% - 0.30% at approximately 45% haematocrit.

The blood meal containing *P. falciparum* NF54 gametocytes was provided in the S3 facility using an artificial membrane feeder system. The mosquitoes were kept at 26°C 80% humidity. At day 11 post infection, infected mosquitoes were sacrificed in 70% ethanol, washed twice in 1 x PBS and dissected in 1% mercurochrome in H_2_O. Oocysts were counted under a light microscope.

### Systemic bacterial injections

One colony of *Escherichia coli* or *Staphylococcus aureus* was picked from agar plates and grown overnight in 5 ml Luria Broth (LB) medium at 37℃ and rotation at 250 rpm. In the morning, liquid cultures were centrifuged at 4,000 x g for 5 min and washed twice in 1 x PBS. Optical density (OD) was measured at 600 nm in 1:1 dilution and concentration for mosquito injections was adjusted with PBS to OD_600_=0.1 for *E. coli* and OD_600_=0.05 for *S. aureus*. Female mosquitoes were anesthetized on ice and injected into the thorax with 69 nl of bacterial mix or 1 x PBS as control using Nanoliter Injector Nanoject II. Injected mosquitoes were transferred to 200 ml paper cups covered with a net, with 10% sucrose cotton pad on top for feeding *ad libitum*, and kept for 4 hours in an insectary at 26℃ and 80% humidity.

### Dissections

Hemocytes were collected by perfusion into a low-binding microcentrifuge tube (Eppendorf) containing 30 *µ*l of insect medium consisting of Schneider’s insect medium, citrate buffer and fetal bovine serum in ratio 6:3:1 as described in^39^. Briefly, 10 mosquitoes were injected into the thorax with 690 nl of the insect medium, using the Nanoliter Injector Nanoject III, and left on ice for 20 min to allow hemocytes to detach from abdominal tissues. Later, a small incision was made with the eye-surgical scissors in the second last abdominal segment. Complete volume of the needle was injected into the animal and the solution with hemocytes was collected with a low-bind pipette tip and placed into a 30 *µ*l of the insect medium in a 2 ml microcentrifuge tube.

Midguts were dissected in a drop of RNase free 1 x PBS by gently pulling off the last abdominal segment, and were divided with a sterile needle at the transition between the thin anterior and thick sac-like posterior midgut.

For the fat body dissection, mosquitoes were pinned to a dissection dish, coated with Steinel hot melt glue, in 50 *µ*l of cold RNase free 1 x PBS. Abdomen was opened along the ventral axis, and its sides were pinned to the dissection dish, exposing the internal organs. Fat body was aspirated using Nanoliter Injector and ejected into a microcentrifuge tube.

Abdominal carcass was separated from the rest of the body, after midgut removal. Carcasses were cleaned from other organs and placed into 2 ml microcentrifuge tube.

### Reverse transcription quantitative polymerase chain reaction (RT-qPCR)

Four hours after systemic infection or PBS challenge, mosquito tissues were dissected as described above. Pools of 10 mosquito tissues were collected per sample for the carcass, midgut and hemocytes. RNAzol (500 µl) was added to each sample, the samples were homogenized with Precellys bead beater at 6,000 x rpm twice for 30 s, with a 30 s break, centrifuged and kept at -80°C until RNA extraction. Total RNA was isolated with DirectZol RNA micro kit, according to manufacturer’s instructions. RNA concentration was measured using Qubit. Total RNA was reverse-transcribed with SuperScript III Kit according to the manufacturer’s instructions.

Mosquitoes from the following conditions: unfed, blood-fed 24 h and *P. falciparum* infected 24 h were collected in 500 *µ*l of Trizol reagent (N=3, n=10), and homogenized with Precellys bead beater (8,000 rpm twice for 30 s, with a 30 s break). DNA was digested using the RapidOut DNA Removal Kit. Total RNA was reverse-transcribed in a volume of 100 *µ*l with 25 *µ*l of dNTPs (2 mM), 0.625 *µ*l of RevertAid H Minus Reverse Transcriptase (200 U/*µ*l), 20 *µ*l of RevertAid H Minus Reverse Transcriptase Buffer (5x), 2.5 *µ*l of Hexamer Primers 100 (*µ*M), 2 *µ*l of Ribolock (20 U/*µ*l) and 2 *µ*g of RNA. Quantitative PCR was performed in a 15 *µ*l volume, containing 7.5 *µ*l FAST SYBR green master mix, 3 *µ*l cDNA template, and 4.5 *µ*l H2O containing forward and reverse primers (Table 1) and 3 *µ*l of cDNA template with a concentration of 0.2 ng/*µ*l of cDNA.

**Table 1:**
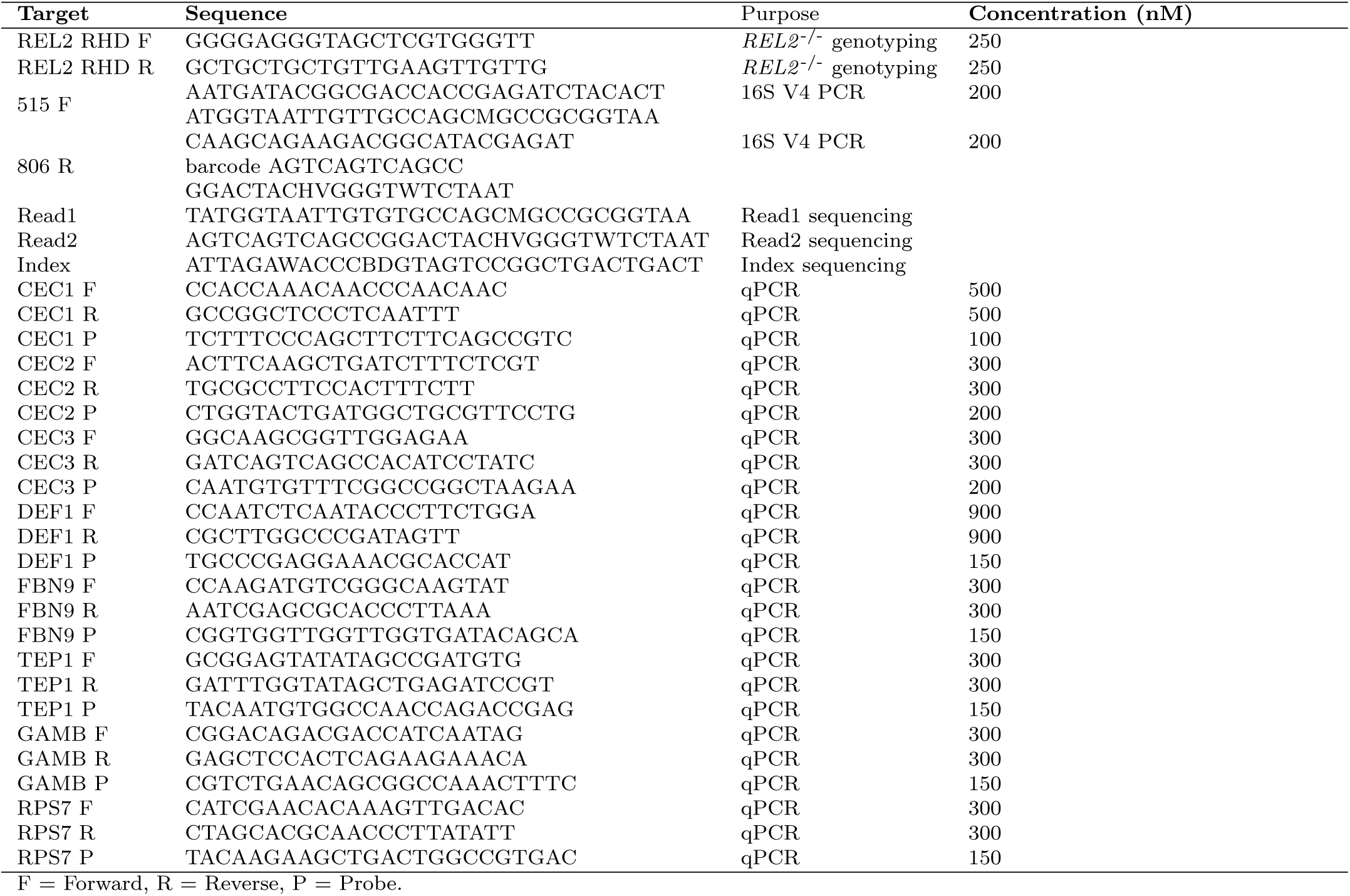
Oligonucleotides.

### RNA-sequencing

Pools of 5 mosquito tissues were collected per sample for the fat body and midgut sections, and hemocytes were extracted from 10 mosquitoes. RNAzol (500 *µ*l) was added and the samples were homogenized with Precellys bead beater at 6,000 x rpm twice for 30 s, with a 30 s break, centrifuged and kept at -80°C until RNA extraction. Total RNA was isolated with DirectZol RNA micro kit and RNA concentration was measured using Qubit. Due to limited RNA quantities for hemocyte samples, cDNA library preparation was performed using a modified protocol for single-cell sequencing with 20 ng of RNA^70^ First strand was reverse-transcribed using oligo-*_dT_* 30 primers. Template Switching Oligos (TSO) were attached to the 5’-end during first strand synthesis. Preamplification was performed for 10 cycles. Quality control was done with the NGS High Sensitivity kit DNF-474-33 (Agilent Fragment Analyzer). Libraries were prepared with Nextera XT DNA Library Preparation Kit. Library enrichment PCR was done with 10 cycles of amplification^70^. Following sample processing and quality evaluation, successful cDNA amplification was achieved for all but one hemocyte (*REL2* ^-/-^ UF) and one anterior midgut sample (WT UF).

Sequencing was performed using NextSeq 150 bp pair-end protocol. Alignment of the reads to reference genome AgamP4.11 was done with bwa mem aligner by EMBL sequencing facility. FastQC program was used for quality control^71^. Gene annotation was performed with *GenomicFeatures* ^72^ using the Anopheles-gambiae-PEST BASEFEATURES AgamP4.12.gtf.gz annotation file, obtained from VectorBase^73^. Count matrix was obtained with *GenomicAlignments* package^72^. The sequenced samples generated approximately 4x10^6^ reads each, with the exception of the three samples of *P. falciparum*-infected mosquitoes, one anterior midgut (*REL2* ^-/-^) and two hemocyte samples (WT and *REL2* ^-/-^) that were excluded from analyses due to low quality and read count (*<* 300,000). For the samples that had passed quality control, transcriptome analyses identified around 10,000 genes per sample.

### RNA-sequencing data analysis

Uniform Manifold Approximation and Projection for Dimension Reduction (UMAP) was performed with *seurat* package^74^. Genes that had sum of counts *<*10 were removed from the count matrix. For creating the seurat object, and clustering, following parameters were used: min.cells=3, min.features=200, k.parameter=8, resolution=0.5, dimensions=14.

Genes that were enriched in each tissue were extracted. Filtering was performed based on following parameters: logfc.threshold =0.25, *P_adj_<*0.05, min.pct=0.25. For plotting the heat plots with gene-marker hits, normalized counts (expressed in natural logarithm) for each tissue were extracted, mean expression for each gene was calculated per strain, tissue and condition, and plotted with *ComplexHeatmap* package in R^75^.

Differential gene expression was analysed on individual tissues with *DESeq2* ^76^. Gene counts were normalized by applying sample-specific size factors to each sample, determined by median ratio method. The counts were transformed with variance stabilizing transformation with *DESeq2*, and Principal Component Analysis (PCA) was performed for top 500 variable genes. Differentially expressed genes (DEGs) were extracted for the following conditions for both anterior and posterior midgut: Unfed (UF) WT vs UF *REL2* ^-/-^, Blood-fed (BF) WT vs BF *REL2* ^-/-^, *Plasmodium*-infected (PF) WT vs PF *REL2* ^-/-^. Afterwards, normalized counts, extracted from the *DESeq2* object were used to compare the significance of the PF DEGs, by comparing their expression with the BF *REL2* ^-/-^ condition.

For Gene Ontology (GO) enrichment analysis, a custom *A. gambiae* database with GO terms was constructed with VectorBase annotations, and differentially expressed genes were screened for enrichment. Statistical significance of identified GO terms (*P*-value) was corrected for multiple-testing using Benjamini-Hochberg correction (*GOATOOLS*, *P_adj_<*0.05)^77^.

### Establishment of a Defensin::GFP/*REL2* ^-/-^ transgenic line for imaging of the Defensin reporter in dissected midgut tissues

*REL2* ^-/-^ mutants were crossed with transgenic mosquitoes expressing the fluorescence reporter *Defensin::GFP* ^41^. The F_2_ generation was COPAS sorted^78^ to establish lines homozygous for the reporter gene, identified by DsRed fluorescence in the eyes, and heterozygous for the *REL2* mutation, marked by GFP in the eyes. In subsequent generations, another round of COPAS sorting based on GFP fluorescence was conducted to produce mixed progenies of 50% control + 50% *REL2* ^-/-^ homozygous mosquito larvae, all homozygous for the reporter construct. The resulting adult mosquitoes were anesthetized on ice and sorted into *REL2* ^-/-^ mutants and WT based on GFP fluorescence in their eyes. The midguts of the selected mosquitoes were dissected in PBS and live-imaged under a Zeiss SMZ18 fluorescence binocular microscope, to reveal the GFP fluorescence patterns of the *Defensin* reporter.

### Mosquito survival

Kaplan-Meier estimate of mosquito survival was calculated over seven to eleven days and plotted in *R* with *survival* and *survminer* incorporated in *ggquickeda* package^79,80^.

### Antibiotic treatment

Control female mosquitoes were given normal 10% sucrose, while antibiotic treatment groups were given sucrose pads supplemented with 75 *µ*g/mL gentamicin sulfate and 100 U/ml penicillin/streptomycin mix, for feeding *ad libitum* from three days after emergence until blood-feeding.

### Quantification of bacterial loads

Mosquitoes were surface-sterilized in 70% ethanol for 30 s and washed in 1 x PBS three times. Individual mosquitoes (n=10 per strain and condition) were homogenized in 100 *µ*l of 1 x PBS. Serial dilutions were prepared and 50 *µ*l were plated on LB agar plates. The plates were kept at 37°C overnight and the colony forming units were counted the next day.

### 16S rDNA amplicon sequencing

Midgut dissections were performed in sterile conditions. Mosquitoes were sacrificed 24 h after blood feeding, collectively washed twice in 70% ethanol for 30 s and three times in sterile 1 x PBS, and three times individually in 1 x PBS. Forceps were cleaned in soapy water and 70% ethanol and sterilized by the Bunsen flame between each sample. Dissected midguts were homogenized with 100 *µ*l of TE buffer and sterile sand with Precellys bead beater at 10,000 rpm twice for 30 s, with a 30 s break. Proteinase K 20 mg/ml (20 *µ*l) in lysis buffer (80 *µ*l) was added to each homogenate and left over-night for digestion at RT. DNA was extracted with NucleoMag VET kit for bacterial and viral DNA, according to the protocol.

Variable region 4 (V4) of *16S* rDNA gene was amplified in triplicates in 25 *µ*l, with 10 *µ*l of 5 Prime HotMasterMix. Barcoded primers with Illumina adapters were used^81^ (Table 1) with 1 *µ*l of DNA template per replicate. The amplicon concentration was assessed with the Agilent Fragment Analyzer and 240 ng of each sample was sequenced paired-end, at the Max Planck Sequencing Facility in Cologne on Illumina HiSeq 2,500 instrument.

### Sequencing data analysis

*16S* rDNA amplicon sequences were processed by *dada2* package^82^ in R. Forward reads were trimmed at 240 and reverse at 200 bp. Silva database, (v. 138) was used to assign species taxonomy to the amplicon sequencing variants (ASV)^83,84^. The resulting ASV count table, taxonomy and metadata were uploaded into a *phyloseq* object^85^. Samples with less than 30,000 reads and taxa with unknown phyla were removed. R *decontam* package was used to remove contaminant taxa^86^. These were classified based on frequency they appeared in the low- versus high-concentration samples.

Alpha diversity was analyzed on non-transformed data. The count matrix filtering based on abundance and prevalence, retained 179 ASVs for further analysis. The data was converted to compositional with *microbiome* package^87^, and agglomerated on the genus level for abundance plotting. Principal component analysis (PCA) was performed with microViz package^88^ after centered-log (crl) transformation of the data^89,90^. Pearson’s correlation coefficients between the 30 most frequent taxa were calculated and networks inferred with *netCoMi* package^91^. The union of the two networks was taken as the layout to facilitate the comparison. The size of the nodes corresponds to the eigenvector values of the taxa in each network. The abundance values were used to assign the colors (viridis paletter). For this purpose, the mean of clr counts was calculated for each genus in both networks, the viridis palette colors were assigned to each value, after which the coloring was manually performed in Inkscape^92^.

### Colonization of mosquitoes with *Serratia*

WT and *REL2* ^-/-^ larvae were reared separately and the pupae (n=300) were cleaned in 500 ml of sterile water in 100 *µ*m cell strainer. All equipment was sterilized by autoclaving or UV-light. Adults were given 10% sterile sugar pads with antibiotic mix containing penicillin/streptomycin (100 U/ml) and gentamicin (50 mg/ml) for two days, followed by sterile sugar for one day. Control group (AB control) continued feeding on 3% sugar, while *Serratia* group was given 3% sugar with *Serratia* spp. cultures grown at 26°C (OD_600_=0.5, corresponding to 1,5x10^8^ bacteria cells). Sterile- or *Serratia*-inoculated sugar pads were given to mosquitoes to feed *ad libitum* for two days. Sugar pads were changed daily. For CFU counting, 3-5 mosquitoes after antibiotic treatment and before blood feeding were homogenized in 100 *µ*l PBS at 6,000 rpm for 10 s. Two serial dilutions were plated (50 *µ*l) on LB agar plates. *Serratia* sp. Ag2 strain, isolated from *Anopheles* mosquitoes by Jiannong Xu, and obtained from BEI resources^44^ was used for colonization.

## Resource availability

### Lead contact

Requests for further information and resources should be directed to and will be fulfilled by the lead contact, Elena A. Levashina

### Materials availability

Mosquito lines generated in this study will be made available upon request.

### Data and code availability

All the original data, as well as raw and processed 16S and RNA-sequencing data, and related code for analysis and visualization have been deposited to Edmond repository https://doi.org/10.17617/3.WSRVSR, and will be publicly available as of the date of publication.

## Acknowledgments

The authors thank the EMBL sequencing facility and Dr. Vladimir Benes for performing the RNA-sequencing and Max Planck Sequencing Facility in Cologne for the 16S rDNA sequencing. We also thank technical assistants of the Vector Biology Department: Daniel Eyermann and Manuela Andres for making the *P. falciparum* cultures, Liane Spohr and Manuela Andres for mosquito infections, and Cornelia Kreschel for the help with mosquito breeding. We are grateful to Marly Erazo for the discussions regarding bacterial colonization, and to Drs. Giulia Costa and Tisheng Shan for critical reading of the manuscript. Finally, we thank current and past Vector Biology lab members for constructive discussions. S.Z.’ work was supported by a Deutsche Forschungsgemeinschaft PhD fellowship (DFG-GRK2046). The project was supported in part by the collaborative CNRS grant “International Research Laboratory” (IRL) 2012-2017.

## Author contributions

E.A.L. Conceptualization and Funding Acquisition. C.K. and E.M. Animal Resources. S.Z., G.E.R., R.G. and C.G.M. Investigation. S.Z. and G.E.R Data Curation. S.Z. Formal Analysis and Visualization. E.A.L. and S.Z. Project Administration, Supervision and Writing.

## Declaration of interest

The authors declare no competing interests.

## Declaration of generative AI and AI-assisted technologies in the writing process

During the preparation of this work the authors used OpenAI chatGPT 4.0 in order to improve the readability, grammar and language of the manuscript. After using this tool, the authors reviewed and edited the content as needed and take full responsibility for the content of the published article.

## Supporting Information

**Figure S1:**
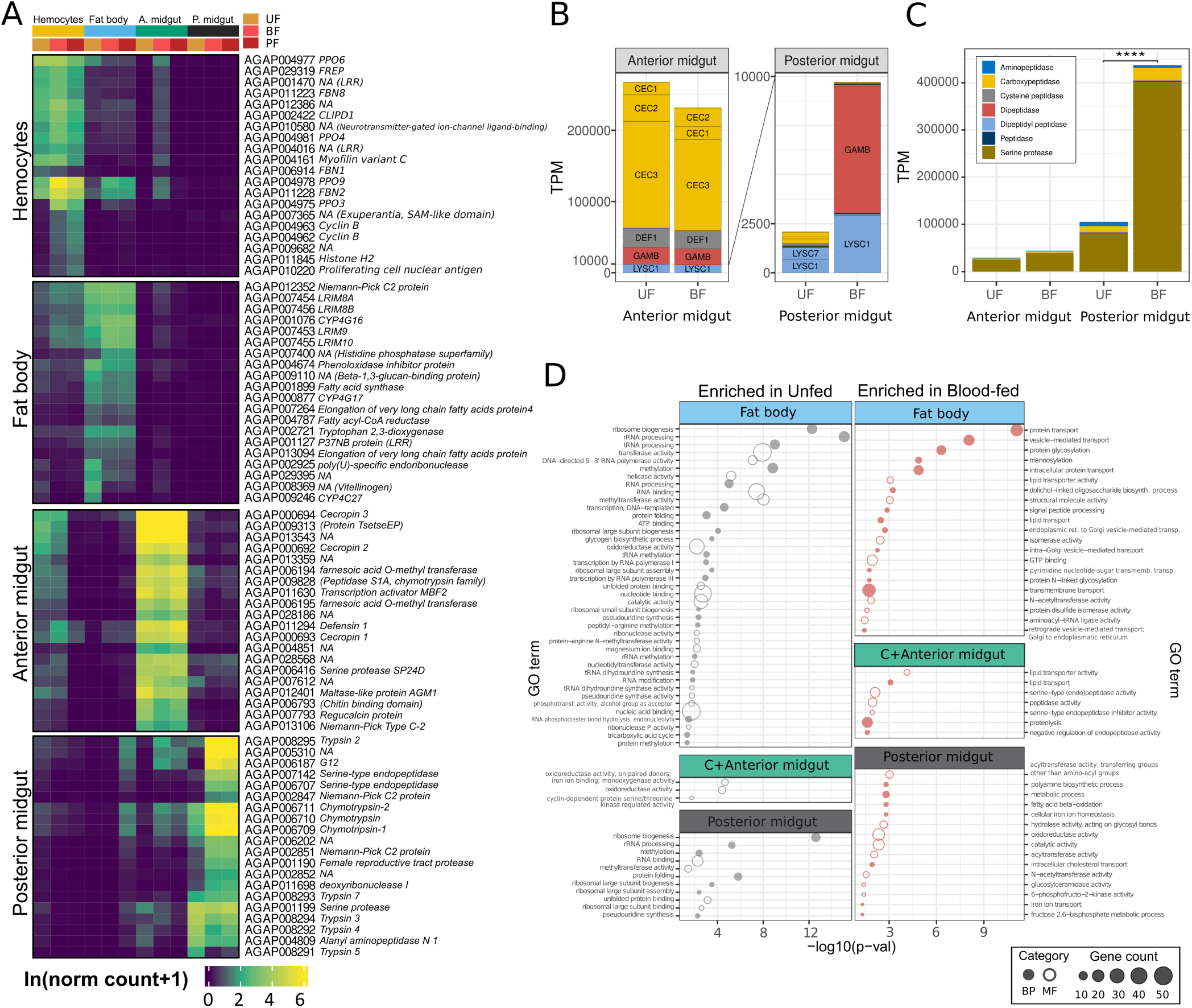
Tissue-specific transcriptional signatures of wild-type mosquitoes. (A) Heatmap displaying the top 20 tissue marker genes identified by UMAP analysis performed in seurat, *P_adj_ <* 0.05. Abbreviations: UF - unfed; BF - blood-fed; PF - *P. falciparum* infected; PPO - prophenoloxydase; FBN - Fibrinogen-like protein; LRIM - Leucine-rich repeat immune protein; CLIP - Clip-domain serine protease; CYP - Cytochrome P450. Each square represents the mean expression from independent experiments (N=3 for all, except N=2 for anterior midgut UF sample). (B, C) Comparison of the anterior and posterior midgut’s investment in AMP (B) and digestive enzyme genes (C), expressed as transcripts per million (TPM), in the unfed vs blood-fed females. Statistically significant differences were evaluated using the Wilcoxon Rank Sum test (***, *p <* 0.0001). (D) Gene ontology (GO) enrichment analysis of tissue-specific responses to blood-feeding (BF). Differentially expressed genes (DEGs) upon BF were compared to the entire *A. gambiae* transcriptome, and GO enrichment was estimated using *GOATOOLS* with *P_adj_ <* 0.05. The results are plotted as -log10 on the x-axis, with bubble size corresponding to the gene count assigned to each GO term.

**Figure S2:**
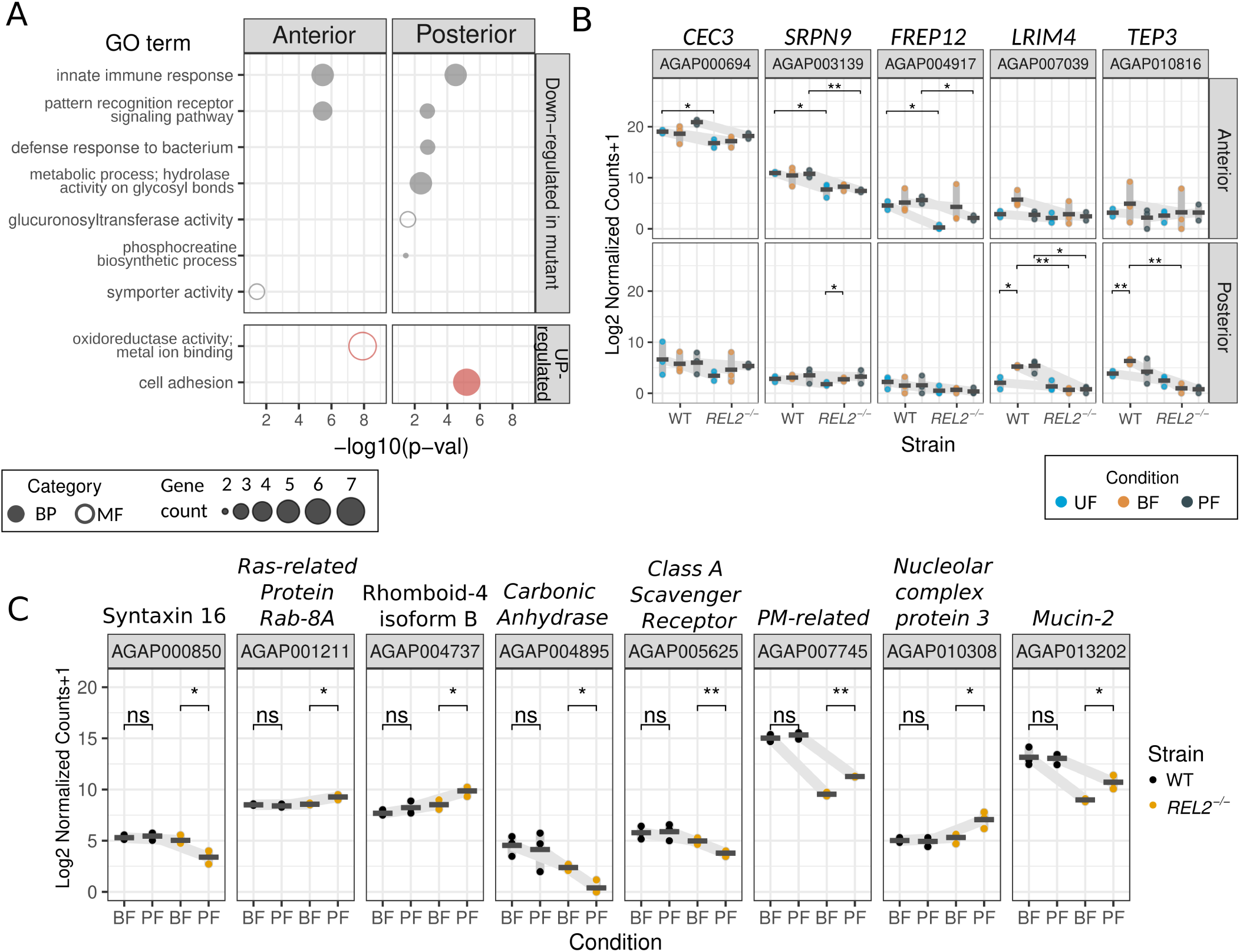
Processes and genes regulated by the REL2 pathway in the mosquito midgut. (A) Gene ontology (GO) processes in the anterior and posterior midgut affected by REL2 deficiency are shown. Differentially expressed genes (down-regulated or up-regulated) in *REL2* ^-/-^ mutants and the wild-type mosquitoes were identified using *DESeq2*. These were then compared to the entire *A. gambiae* transcriptome to identify enriched GO terms using *GOATOOLS*, with *P_adj_ <* 0.05. The results are plotted as -log10 on the x-axis. Bubble size corresponds to the gene count assigned to each GO term. (B) Genes regulated by the REL2 pathway in either anterior or posterior midgut were identified using *DESeq2*. Statistical significance was evaluated using a Student’s t-test (*, *p <* 0.05; **, *p <* 0.01; ***, p *<* 0.001; ****, p *<* 0.0001). UF: unfed; BF: blood-fed; PF: *P. falciparum* infected. Dots represent independent experiments. Mean expression levels (indicated by crossbars) for each line are grouped by experimental conditions and connected by gray lines. (C) Differentially expressed genes (DEGs) whose expression is regulated by the REL2 pathway after *P. falciparum* infection (PF). DEGs were identified using *DESeq2* analysis and their normalized counts were compared to those in blood-fed (BF) samples as the baseline. Statistically significant differences in expression were assessed using a Student’s t-test (ns *>* 0.05; *, *p <* 0.05; **, *p <* 0.01).

**Figure S3:**
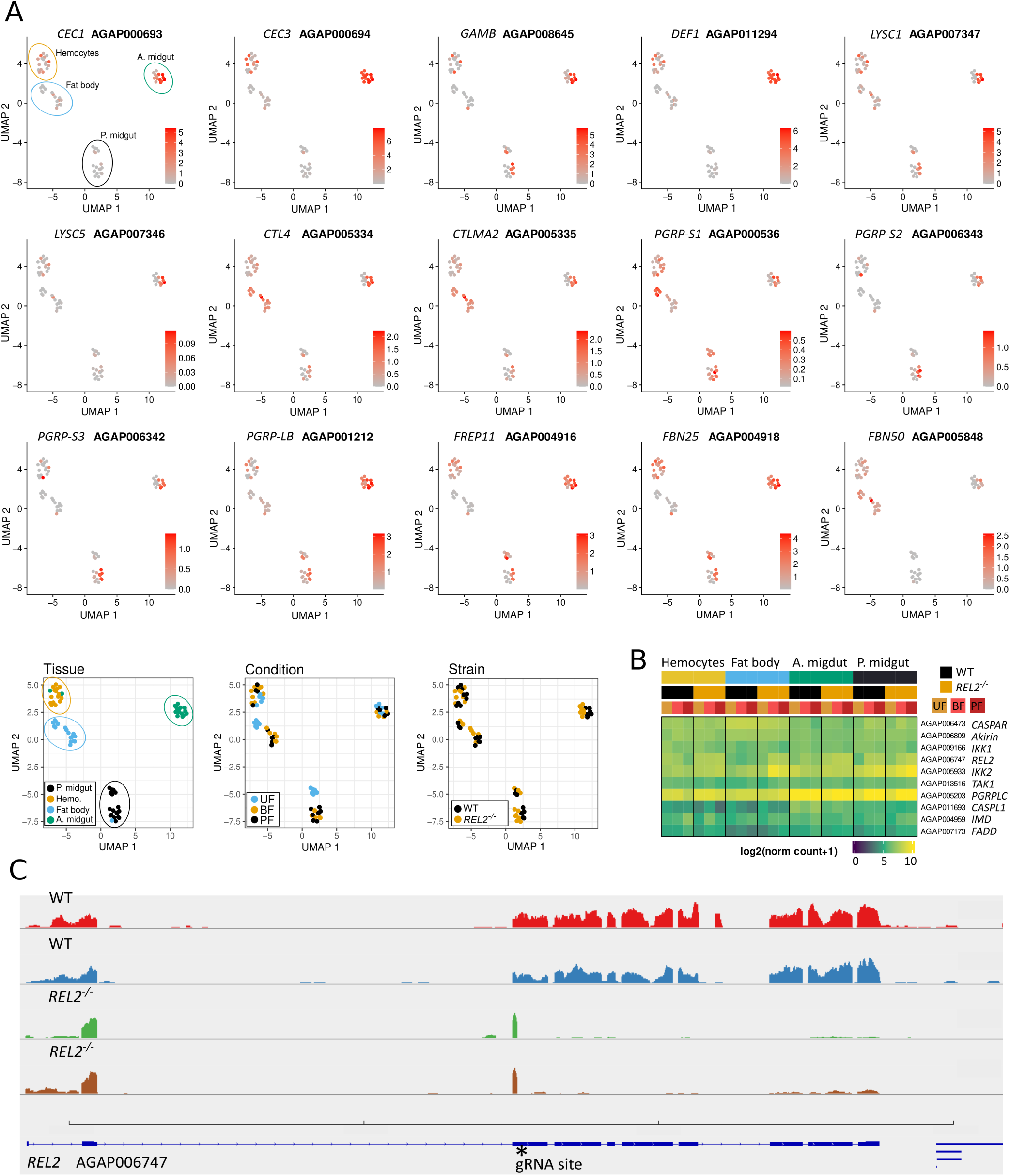
Tissue-specific expression of REL2 targets and pathway components. (A) The expression levels of REL2 target genes are depicted using UMAP dimension reduction plots (rows 1-3). Each dot represents a single sample. Clusters identified by UMAP analysis (row 4) are colored by tissue, condition and mosquito line for easier comparison. (B) Heatmap illustrates the expression levels of genes encoding REL2 pathway components. Each square represents the mean of three independent biological experiments. (C) Alignment of RNA-seq reads to the *REL2* locus. The first two rows show transcript coverage of the *REL2* locus in WT mosuitoes, while the second two rows display transcript alignments in *REL2* ^-/-^ mutants. Schematic depiction of exon-intron organization of the *REL2* locus. The gRNA target site where the GFP cassette was inserted is shown with a star. Abbreviations: RHD, REL Homology Domain; ANK, Ankyrin-Repeat Domain.

**Figure S4:**
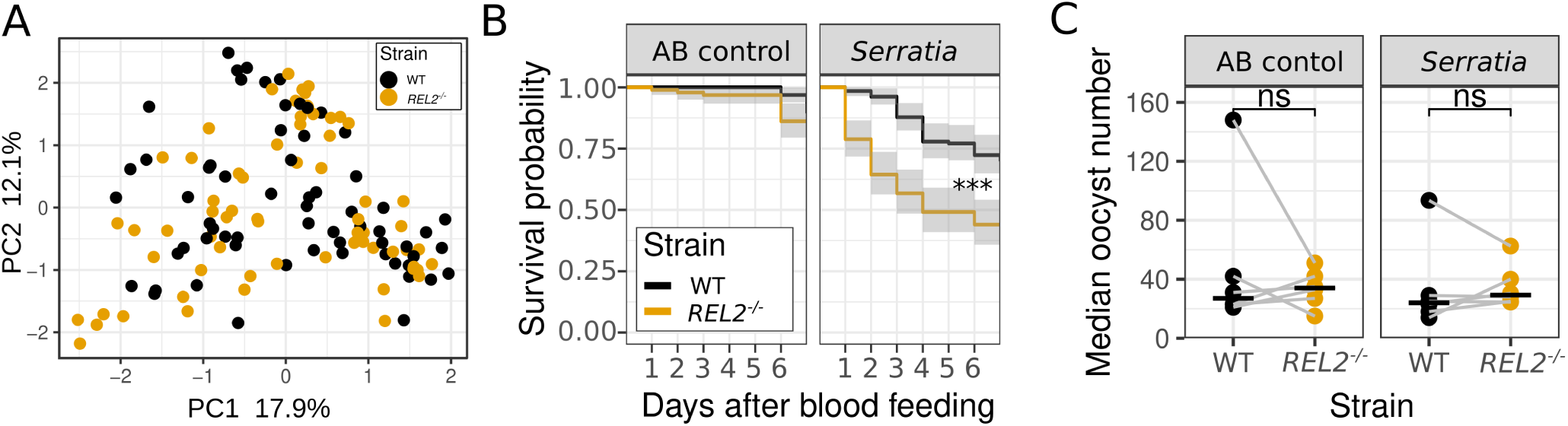
Impact of *Serratia* spp. colonization on mosquito survival rates and *P. falciparum* oocyst loads. (A) PCA of the beta-diversity in mosquito microbiome data, following center-log transformation of read counts. The data is color-coded by mosquito line: WT (black) and *REL2* ^-/-^ (yellow). (B) The effect of *Serratia* spp. colonization on mosquito survival over a period of 7 days post blood-feeding. WT and *REL2* ^-/-^ mosquitoes after eclosion were treated with antibiotics. Mosquitoes were orally colonized with *Serratia* sp. Ag2 bacterial cultures (OD_600_=1) by sugar feeding (*Serratia* group) or left uncolonized (AB control). Survival was assessed using the Kaplan-Meier estimate and statistically significant differences were evaluated with the Log-rank test (***, *p <* 0.0001, N=3, n ≥ 20). (C) Median oocyst counts per experiment for antibiotic-treated mosquitoes (AB control) and mosquitoes colonized with *Serratia* sp. Ag2 (*Serratia*, OD_600_=0.5, N=6, n=12-50). Horizontal bars show medians. Statistically significant differences were evaluated by the Wilcoxon Rank Sum test (ns, *p >* 0.05).

## Notes

### Competing Interest Statement

The authors have declared no competing interest.

